# Estimating the time since admixture from phased and unphased molecular data

**DOI:** 10.1101/2020.09.10.292441

**Authors:** Thijs Janzen, Verónica Miró Pina

## Abstract

After admixture, recombination breaks down genomic blocks of contiguous ancestry. The breakdown of these blocks forms a new ‘molecular clock’, that ticks at a much faster rate than the mutation clock, enabling accurate dating of admixture events in the recent past. However, existing theory on the break down of these blocks, or the accumulation of delineations between blocks, so called ‘junctions’, has mostly been limited to using regularly spaced markers on phased data. Here, we present an extension to the theory of junctions using the Ancestral Recombination Graph that describes the expected number of junctions for any distribution of markers along the genome. Furthermore, we provide a new framework to infer the time since admixture using unphased data. We demonstrate both the phased and unphased methods on simulated data and show that our new extensions performs better than previous methods, especially for smaller population sizes and for more ancient admixture times. Lastly, we demonstrate the applicability of our method on an empirical dataset of labcrosses of yeast (*Saccharomyces cerevisae*) and on two case studies of hybridization in swordtail fish and *Populus* trees.

## 1 Introduction

The traditional view where taxa accumulate incompatibilities over time and gradually become reproductively isolated from each other – speciation – has led to insight into the processes generating and maintaining biodiversity (Coyne and Orr, 2004). However, it has become apparent that reticulate evolution is common: taxa do not necessarily only branch, they can also come back together – hybridisation – (Abbott et al., 2013). This means that, while the ancestry of non-recombining DNA can be traced linearly back in time (e.g. mitochondrial lineages), the ancestry of genomes backward in time will branch at admixture events. In plants, it has been known for quite some time that hybridization is a widespread phenomenon that can not only generate viable offspring, but also potentially lead to the formation of new taxa, and ultimately, species (Grant, 1981). It has long been debated whether this process is also common in animals, but over the past few years numerous examples have appeared, including, but not limited to, butterflies (Mavárez et al., 2006; Capblancq et al., 2015), cichlid fishes (Koblmüller et al., 2007; Keller et al., 2013), warblers (Brelsford et al., 2011), fruit flies (Schwarz et al., 2005) and sculpins (Nolte et al., 2005).

Understanding the timeline of these hybridization events is paramount in obtaining a full understanding of the process and its impact. A ‘recombination clock’ is particularly useful for studying recent evolutionary dynamics since detectable recombination events are markers of admixture (Baird, 2006), and the statistical association across loci created by admixture (admixture linkage disequilibrium) decays slowly (Baird, 2015). After admixture of two taxa, contiguous genomic blocks are broken down by recombination over time. The delineations between these blocks were termed ‘junctions’ by Fisher (1949, 1954), and inheritance of these junctions is similar to that of point-mutations. Further work on the theory of junctions has shown how they accumulate over time for sib-sib mating (Fisher, 1954), self-fertilization (Bennett, 1953), alternate parent-offspring mating (Fisher, 1959; Gale, 1964), a randomly mating population (Stam, 1980; Baird et al., 2003), and for sub-structured populations (Barton, 1983; Baird, 1995; Chapman and Thompson, 2002, 2003).

So far, applying the theory of junctions has proved difficult, as it requires extensive genotyping of the admixed individuals, but also of the source taxa. With the current decrease in genotyping costs (Muir et al., 2016), such analyses are coming within reach, and frameworks are being developed that assist in inferring local ancestry and detecting junctions, given molecular data of source and admixed taxa (Paşaniuc et al., 2009; Maples et al., 2013; Guan, 2014; Corbett-Detig and Nielsen, 2017; Medina et al., 2018; Svedberg et al., 2021). Nevertheless, molecular data always paints an imperfect image of ancestry along the genome, and inferring the number of junctions in a chromosome remains limited by the number of diagnostic markers available (see Fig 1, first panel). Previous work on the theory of junctions does not take into account the effect of a limited number of genetic markers, and so far this effect had to be corrected using simulations (MacLeod et al., 2005; Buerkle and Rieseberg, 2008). Recent work by Janzen et al. (2018) resolves this issue by extending the theory of junctions with the effect of using a limited number of markers, but they had to assume an evenly spacing of markers. However, molecular markers are rarely evenly spaced. The first result we present here is an extension of the theory of junctions which includes the effect of marker spacing on inferring the number of junctions in a genome and that has the advantage of working well with fewer markers.

**Fig 1.**
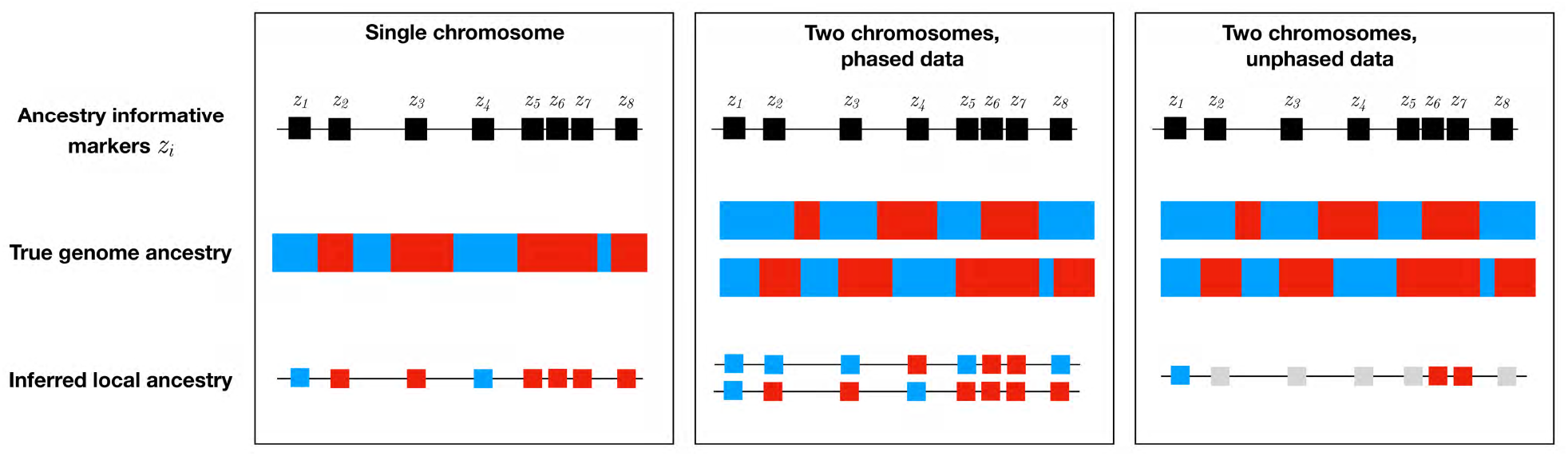
Visual depiction of the observed data. We show the differences between the type of data generated by the three methods we present in this paper. On each panel, the chromosome in the center is colored according to ancestry (blue represents source taxa 𝒫 and red represents source taxa 𝒬). Above the chromosome are indicated the locations of ancestry informative markers *z*_*i*_. Resulting inferred ancestry on these markers is shown below, where grey indicates heterozygous ancestry. The first panel represents the one chromosome method. There are 7 junctions in the chromosome, but only 3 are observed in the data due to a limited marker coverage. Notice that a junction in the inferred ancestry corresponds to an odd number of junctions in the “true” genome ancestry and conversely, that an even number of junctions in the “true” ancestry yields no inferred junctions. The second and third panels represent the methods that use information from two chromosomes. In the second panel data is phased whereas in the second panel data is unphased.

Furthermore, most existing theory on the accumulation of junctions is developed for the case where ancestry can be determined within a single chromosome (with the exception of ancestry hmm, (Corbett-Detig and Nielsen, 2017)). For diploid species, sequencing data presents itself as the pileup of ancestry across both chromosomes, requiring an additional step to separate the contributions of both chromosomes, called ‘phasing’ (see Fig 1, second and third panel). Phasing methods can be classified into three main categories, including haplotype-resolved genome sequencing, pedigree-based methods and statistical methods. Haplotype-resolved genome sequencing methods (reviewed in Snyder et al. (2015)), yield accurate results, but are expensive and require a large number of sequences in order to be able to resolve the required haplotypes (but see (Lutgen et al., 2020)). Pedigree based phasing methods do not require resolved haplotypes, but instead rely on accurate genotyping of related individuals to resolve the required phase. These methods often yield good results but their application has been limited to humans, where large pedigree datasets are available (Browning and Browning, 2011; Loh et al., 2016a; Kong et al., 2008). Statistical methods do not require large pedigree datasets, and are based on recombination rate estimates and allele frequencies in a population. While some of these methods make use of a reference genome (for example Eagle (Loh et al., 2016b), Beagle (Browning and Browning, 2007) or ShapeIt (O’Connell et al., 2016)), others allow *de novo* haplotype aware assembly (for example POLYTE (Baaijens and Schönhuth, 2019)). Further improvements include the usage of third-generation sequencing data (Tourdot and Zhang, 2019; Kronenberg et al., 2019; Ebler et al., 2019; Tangherloni et al., 2019). However, data from hybrid populations is not often available in this form. Across these three groups of methods, the overarching theme is that phasing is often costly and accuracy can be left wanting. Yet, inclusion of information from both chromosomes is expected to improve inference of the onset of admixture considerably and hence expansion of the theory of junctions towards a framework that takes into account data from both chromosomes is warranted.

Here we provide a framework to estimate the time since admixture using phased or unphased data from two homologous chromosomes, taking into account marker spacing along the chromosome. Our framework is based on modelling the joint genealogy of loci that are located in the same chromosome or in two homologous chromosomes, using the Ancestral Recombination Graph (ARG) (Hudson, 1983; Griffiths, 1991; Griffiths and Marjoram, 1997). It has the advantage of being fast since it relies on mathematical computations and does not require simulations. It has been implemented in the R package ‘junctions’. In comparison with previous methods (ancestry hmm, Corbett-Detig and Nielsen (2017)), our method has the advantage of working well with small population sizes and larger admixture times.

Our paper is organised as follows. In section 2, we introduce our model, which is a simplified version of the ARG and present three maximum-likelihood methods to infer the time since admixture in hybrid populations: the first method uses information from a single chromosome, the second method uses phased data from two homologous data and the third method uses unphased data from two homologous chromosomes. At the end of section (2.4), we validate our methods using simulations. In section 3, we provide a detailed comparison to previous methods. Lastly, in section 4 we apply them to a dataset from experimental evolution in yeast and to two case studies of hybridization in swordtail fish and *Populus* trees.

## 2 Materials and methods

### 2.1 Mathematical model

We assume a diploid population that evolves according to Wright-Fisher dynamics, i.e. generations are non-overlapping, mating is random and all individuals are hermaphrodites. We only keep track of one chromosome (or one pair of chromosomes), assuming that the accumulation of junctions on different pairs of chromosomes is independent of each other (see Section 2.2.1). We assume that hybridization occurred *T* generations ago between two source taxa, 𝒫 and 𝒬. The proportion of individuals from source 𝒫 in the initial generation is *p* and the proportion of individuals from source 𝒬 is *q* = 1 − *p*. We define the initial heterogenicity *H*_0_ := 2*pq* as the probability that, when sampling two individuals from this generation they are from different source taxa. This also represents the probability that at any given locus, one allele can be traced to source taxon 𝒫 and the other allele to 𝒬. Following Janzen et al. (2018), instead of referring to the lesser known term of heterogenicity, we use the term of heterozygosity in the manuscript throughout because these two concepts are mathematically equivalent.

We assume that the length of the chromosome is *C* Morgan and that there are *n* molecular markers whose positions are given by (*z*_1_, …, *z*_*n*_) ∈ [0, *C*]. For two consecutive markers at sites *z*_*i*_ and *z*_*i*+1_, we define *d*_*i*_ = *z*_*i*+1_ − *z*_*i*_, the distance between them in Morgan. We assume that there are enough markers on the chromosome such that the *d*_*i*_’s are small compared to 1. The genealogy of these *n* (or 2*n* for a diploid genome) loci is given by the Ancestral Recombination Graph (ARG), defined in Hudson (1983); Griffiths (1991); Griffiths and Marjoram (1997). The ARG is a branching-coalescence process which follows backwards in time the ancestry of loci sampled in the present population (time 0). If two loci belong to the same block at time *t*, they are identical-by-descent (IBD) with respect to generation *t*, i.e. the sampled loci have been inherited from the same individual living *t* units of time ago.

Although the ARG for many loci has complicated transition rates and is a computationally intensive model, here we consider only two loci (or two pairs of loci for a diploid genome) at a time. This is sufficient since for our maximum likelihood approach we only use the expected number of junctions – and not its variance or higher moments (see e.g. equation 3).

We assume that the number of diploid individuals *N* is large, so that we can neglect some transitions (double coalescences and simultaneous coalescence and recombination) and that *d*_*i*_ ≪ 1 so that there is no more than one crossover per generation between two molecular markers and the mutation rates are small enough so that we can neglect mutations that happened between the admixture time and the present.

### 2.2 Two sites, one chromosome

The aim of this section is to derive a formula for the expected number of observed junctions on one chromosome given the population size *N*, the distances between the markers (*d*_1_, …, *d*_*n*_) and the initial heterozygosity *H*_0_ := 2*pq*. We start by considering two consecutive loci *z*_*i*_ and *z*_*i*+1_ sampled in the same chromosome in the present population. The ARG for these two sites has two possible states:

- If the loci are in state (*z*_*i*_ ∼ *z*_*i*+1_) at time *t*, it means that the two sampled loci have been inherited from *the same individual* living *t* generations before, i.e. that they are IBD.
- If the loci are in state (*z*_*i*_ ≁ *z*_*i*+1_), it means that the two loci have been inherited from different individuals.

To model the observed junctions, we look at identity-by-descent with respect to time *T*, when admixture took place. Recall that we only model observed junctions, i.e. where adjacent markers have alleles corresponding to different source taxa (which corresponds to an odd number of “true” junctions, see 1 and its caption).

- If the two loci are in state (*z*_*i*_ ∼ *z*_*i*+1_) at time *T*, the ancestor can be from source 𝒫 (with probability *p*) or red (with probability *q*). In both cases, we observe no junction.
- If the two loci are in state (*z*_*i*_ ≁ *z*_*i*+1_) at time *T* (not IBD), they have two different ancestors at generation *T*, which can be:
  – Both blue (with probability *p*^2^) or both red (with probability *q*^2^), in this case there is no junction.
  – One blue and one red (with probability 2*pq*), in this case, there is a junction in the chromosome sampled in the present.
  The dynamics of the ARG are controlled by two types of events:
- **Recombination** (*z*_*i*_ ∼ *z*_*i*+1_) → (*z*_*i*_ ≁ *z*_*i*+1_) with probability *d*_*i*_,
- **Coalescence** (*z*_*i*_ ≁ *z*_*i*+1_) → (*z*_*i*_ ∼ *z*_*i*+1_) with probability 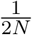.

Since *N* ≫ 1 and the *d*_*i*_ ≪ 1, other events (such as simultaneous coalescence and recombination events) have probabilities that are negligible. This yields the following transition matrix:

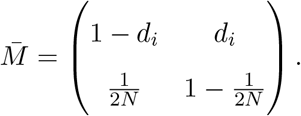

Let 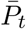 be the probability vector at time *t* for this Markov chain with two states. 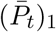 is the probability of (*z*_*i*_ ∼ *z*_*i*+1_) at time *t* and 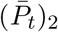 the probability of (*z*_*i*_ ≁ *z*_*i*+1_) at time *t*. We have 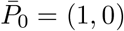 (in the present we sample the two loci in the same individual) and 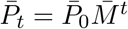. We denote by 𝕡 (*J*_*T*_ (*z*_*i*_, *z*_*i*+1_)) the probability that a junction is observed between *z*_*i*_ and *z*_*i*+1_ (conditioning on the fact that hybridization occurred *T* generations ago). We have

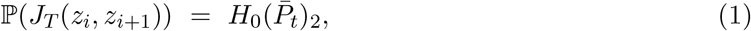

which corresponds to the probability that the two loci were carried by different individuals *T* generations ago and they from different source taxa (see Fig 2, left panel).

**Fig 2.**
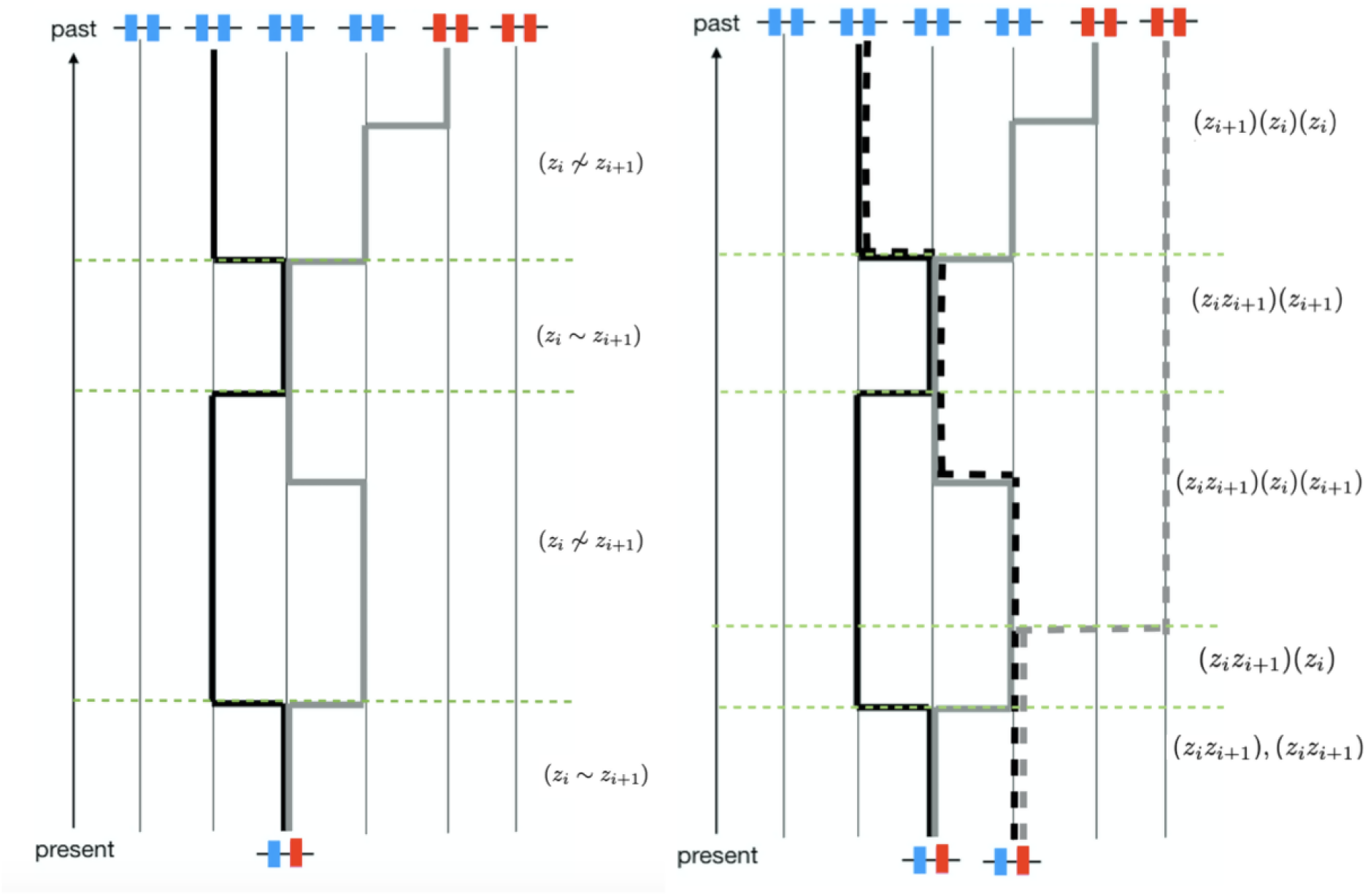
The ARG with two markers. Each color represents one source taxa (𝒫 and 𝒬). The black and grey lines (or dotted lines) represent the ancestral lineage of each marker. In the left panel, we show the ARG for two markers in one chromosome. In the present, there is an observed junction between the two markers. In the past (*t* generations ago, when hybridization took place), each lineage is carried by a different individual and these two individuals are from different source taxa. The right panel shows the ARG for two markers in two homologous chromosomes.

Solving equation (1) gives

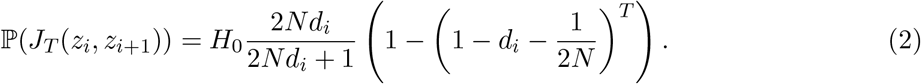

Let 𝔼 (*J*_*T*_) be the expected number of observed junctions on one chromosome, we have

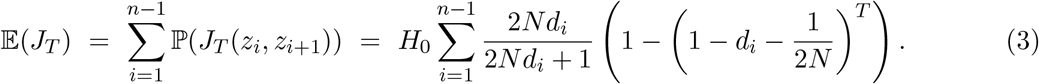

#### 2.2.1 Maximum Likelihood

For each chromosome, we can calculate the likelihood of observing the data, where the data are *n* − 1 pairs of markers (*z*_*i*_ ∼ *z*_*i*+1_), which is given by

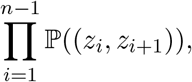

where 𝕡((*z*_*i*_, *z*_*i*+1_)) is the probability of observing the pair of markers (*z*_*i*_, *z*_*i*+1_) given by:

- 𝕡((*z*_*i*_, *z*_*i*+1_) = 𝕡(*J*_*T*_ (*z*_*i*_, *z*_*i*+1_)), if there is a junction between the two markers. This is given by equation (2).
- 𝕡((*z*_*i*_, *z*_*i*+1_) = 1 − 𝕡(*J*_*T*_ (*z*_*i*_, *z*_*i*+1_)), if there is no junction between the two markers.

For different chromosomes sampled in the same individual, we can assume independence between chromosomes and the full likelihood of observing the data, given the time since admixture, is given by the product of these likelihoods, e.g.:

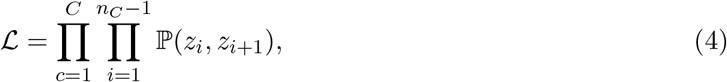

where *C* indicates the number of chromosomes and *n*_*C*_ indicates the number of markers on chromosome *C*.

When we have data from several individuals (which are not independent, due to coalescence), we first compute the maximum likelihood estimator of *T* for each individual (using (4)) and then we average across all individuals in the sample. Confidence intervals (CI’s) reported are then the 95 % interquartile range in the age estimates across all individuals.

**Table 1.** States of the reduced ARG. *n*_*i*_ (resp. *n*_*i*+1_) denotes the number of ancestors of site *z*_*i*_ (resp. *z*_*i*+1_) and *n*_*tot*_ the total number of ancestors to the sample.

| State | | $n_i$ | $n_{i+1}$ | $n_{tot}$ |
| --- | --- | --- | --- | --- |
| $S^1$ | $(z_i z_{i+1}), (z_i z_{i+1})$ | 2 | 2 | 2 |
| $S^2$ | $(z_i z_{i+1})(z_i)(z_{i+1})$ | 2 | 2 | 3 |
| $S^3$ | $(z_i)(z_i)(z_{i+1})(z_{i+1})$ | 2 | 2 | 4 |
| $S^4$ | $(z_i z_{i+1})(z_i)$ or $(z_i z_{i+1})(z_{i+1})$ | 2 (or 1) | 1 (or 2) | 2 |
| $S^5$ | $(z_i)(z_{i+1})(z_{i+1})$ or $(z_{i+1})(z_i)(z_i)$ | 1 (or 2) | 2 (or 1) | 3 |
| $S^6$ | $(z_i z_{i+1})$ | 1 | 1 | 1 |
| $S^7$ | $(z_i), (z_{i+1})$ | 1 | 1 | 2 |

### 2.3 Two sites, two chromosomes

We consider two consecutive loci *z*_*i*_ and *z*_*i*+1_, which are at distance *d*_*i*_ (in Morgan), that we sample in two homologous chromosomes. The ARG for these 2 sites in 2 chromosomes has 7 states (see Durrett (2008), Chapter 3). To describe them, we borrow the notation from Durrett (2008) and we write (*z*_*i*_*z*_*i*+1_) to indicate that sites *z*_*i*_ and *z*_*i*+1_ are IBD, and notation (*z*_*i*_) (or (*z*_*i*+1_)) to indicate that the ancestor to *z*_*i*_ (or (*z*_*i*+1_)) is only ancestor to one of the two sites. The resulting 7 states are summarized in Fig 1. An example of realization of this process is shown in Fig 2 (right panel).

The initial state is *S*^1^ because in the present time we sample two different loci in two different chromosomes. The transition matrix of the ARG with 2 loci and a sample size 2 can be approximated, when *N* ≫ 1 by

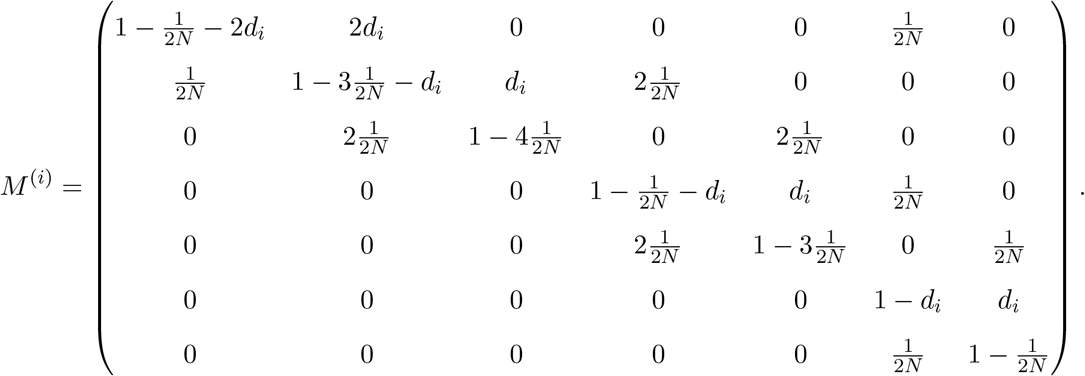

All other potential events (e.g. double crossovers or simultaneous crossover and coalescence events) have probabilities that are negligible compared to 1*/*2*N*.

Let 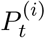 be the vector containing the probabilities of observing each of the states (*S*^1^, …, *S*^7^) at time *t*. 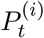 satisfies

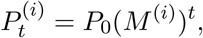

where *P*_0_ = (1, 0, 0, 0, 0, 0, 0), since at time 0 we sample all loci in two homologous chromosomes. This equation can only be solved numerically. Recall that the stationary distribution of this process *P*^(*i*)^ satisfies

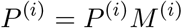

and has the analytical expression

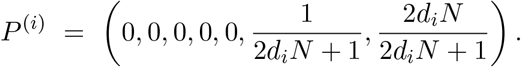

Thus, for large values of *t* the system reduces to states *S*^6^ and *S*^7^, which means that each locus has only one ancestor i.e. forwards in time the process has reached fixation (at each locus). Recall that state *S*^6^ is the state where there is one ancestor for the sample thus we observe no junctions on either chromosome. Furthermore, recall that state *S*^7^ is the state where there are two ancestors, one for the first locus and one for the second locus, and with probability 2*pq* each one of them comes from a different source taxa. This is exactly the probability of observing a junction when *t* → ∞ for one chromosome (equation 2). In other words, when *t* is very large, fixation is reached and the two sampled chromosomes are homozygous so the problem reduces to the single chromosome case.

#### 2.3.1 Maximum Likelihood, phased data

We first consider the case of phased data. Each pair of homologous markers can be in one of four states:

- *PP* i.e. both homologous markers carry the allele from source 𝒫,
- *QQ* i.e. both homologous markers carry the allele from source 𝒬,
- *PQ* i.e. the marker on the first chromosome carries the allele from source 𝒫 and the marker on the second chromosome carries the allele from source 𝒬,
- *QP* i.e. the marker on the first chromosome carries the allele from source 𝒬 and the marker on the second chromosome carries the allele from source 𝒫.

The data can then be represented as a sequence (*O*_*i*_, 1 ≤ *i* ≤ *n*) that takes values in {*PP, QQ, PQ, QP*} such that *O*_*i*_ is the state of the i-th marker. To derive a maximum likelihood formula for the time since admixture *T*, we compute the probability of each sequence in {*PP, QQ, PQ, QP*}*n* given *T, N, C*, the distances between the *n* loci and the initial heterozygosity *H*_0_.

We want to compute the probability of our observations (*O*_1_, …, *O*_*n*_). These *n* observations are not independent, as there are non-trivial correlations between loci along the chromosome. However, we can neglect long-range dependencies and assume that *O*_*i*_ only depends on *O*_*i*−1_, i.e. that the probability of observing (*O*_1_, …, *O*_*n*_), *t* units of time after hybridization is

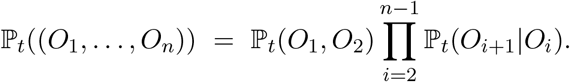

Recall that ignoring long-range dependencies is a natural approximation and it has been used for example by McVean and Cardin (2005) to define the sequentially Markov coalescent. To compute ℙ_*t*_(*O*_*i*+1_|*O*_*i*_), we use the ARG for markers at *z*_*i*_ and *z*_*i*+1_ denoted by 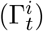 (and to compute ℙ (*O*_1_, *O*_2_), we use 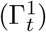)). For example, we can observe *O*_1_ = *PP* and *O*_2_ = *QQ* if:

- 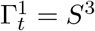 and the two ancestors for locus 1 are from source taxa 𝒫 and the two ancestors for locus 2 from 𝒬, which happens with probability *p*^2^*q*^2^ or,
- 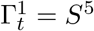, with probability 1*/*2 there are two ancestors for locus 1 and one for locus 2 and they are from desired source taxa with probability *p*^2^*q*. With probability 1*/*2 there is one ancestor for locus 1 and two for locus 2 and they are from the desired source taxa with probability *pq*^2^ or,
- 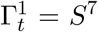 and the ancestor to 1 is from source taxa 𝒫 and the ancestor to 2 from 𝒬, which happens with probability *pq*.

To sum up, when *O*_1_ = *PP* and *O*_2_ = *QQ*,

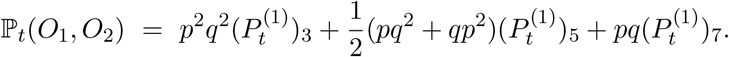

The probabilities for all combinations of *O*_1_ and *O*_2_ are listed in Fig 3. To compute P_*t*_(*O*_*i*+1_|*O*_*i*_) we use Bayes’ formula:

**Fig 3.**
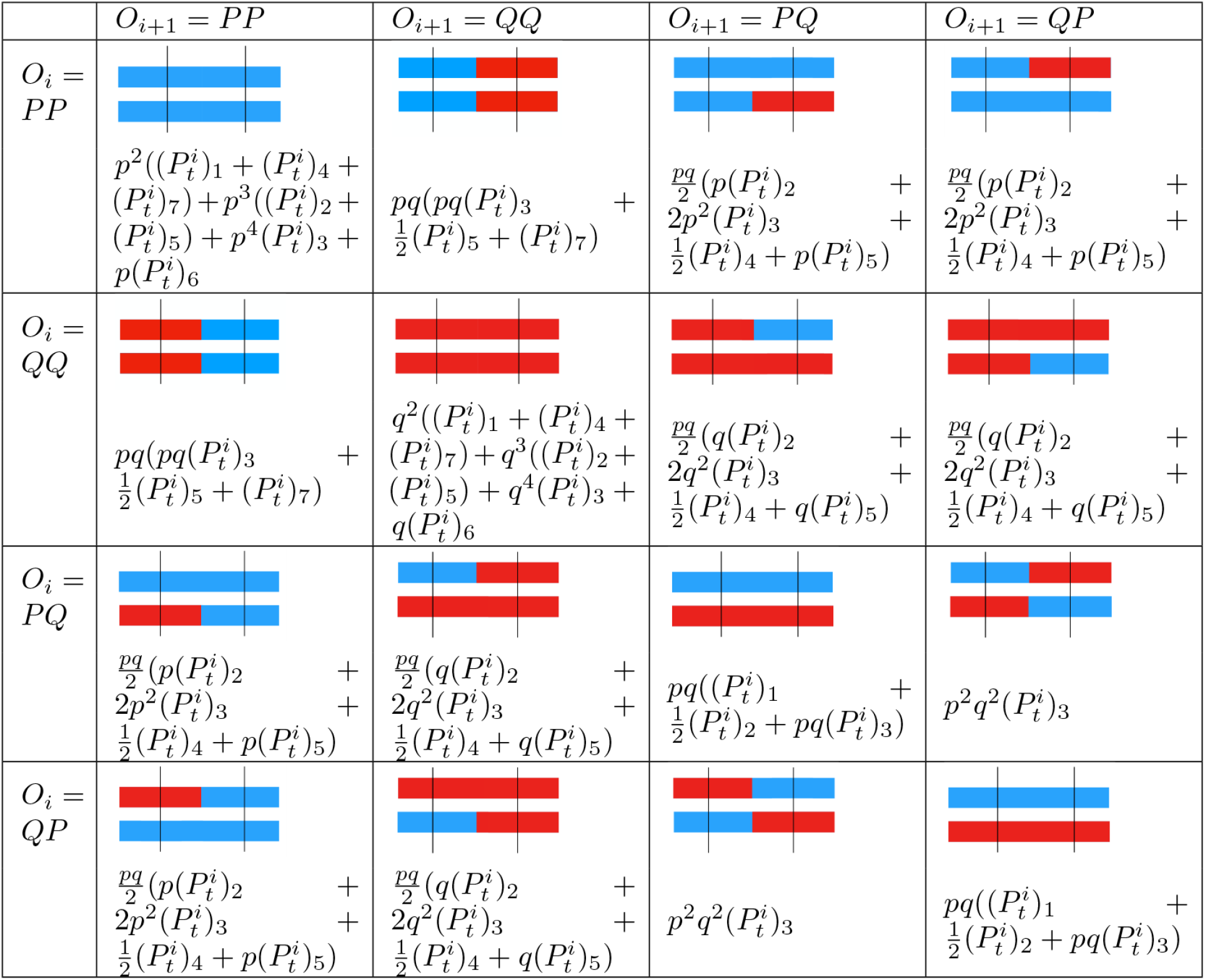
ℙ_*t*_(*O*_*i*_, *O*_*i*+1_) for phased data. The allele from source taxa 𝒫 is represented in blue and the allele from source taxa 𝒬 is represented in red.

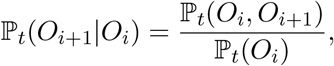

where, using the total probability theorem, ℙ_*t*_(*O*_*i*_) can be obtained by summing over the appropriate row in Fig 3. Then, the total probability of observing the data, given *N* and *t*, i.e.

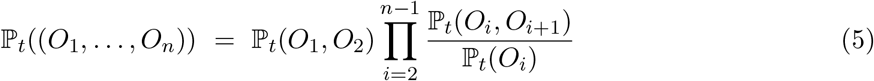

can be maximized in order to find the maximum likelihood estimator of *t* and *N*. If the data consists of multiple chromosomes from the same individual, we can calculate the joint likelihood of observing the data, given *t* and *N* by assuming independence across chromosomes and calculating the likelihood as the product across chromosomes. As in the one chromosome case, we provide estimates of *T* by averaging across the different individuals of the sample, and the CI’s reflect the 95 % interquartile range of the *T* estimates across the different individuals of the sample.

#### 2.3.2 Maximum Likelihood, unphased data

If the data is unphased, we cannot distinguish which allele is in which of the two homologous chromosomes. We can observe one of these three states at each marker:

- *P* i.e. we only observe the allele from source 𝒫, i.e. both chromosomes carry the allele from source 𝒫,
- *Q* i.e. we only observe the allele from source 𝒬.
- *x* i.e. we observe both alleles, i.e. each one of the two homologous chromosomes carries a different allele.

The data can then be represented as a sequence (*O*_*i*_) of length *n* that takes values in *{P, Q, x}* such that *O*_*i*_ is the state of the i-th marker. We can perform exactly the same method, as in the previous section, except that now the probabilities of each state are given by Fig 4. Again, if the data consists of multiple chromosomes from the same individual, we can calculate the joint likelihood of observing the data, given *t* and *N* by assuming independence across chromosomes and calculating the likelihood as the product across chromosomes.

**Fig 4.**
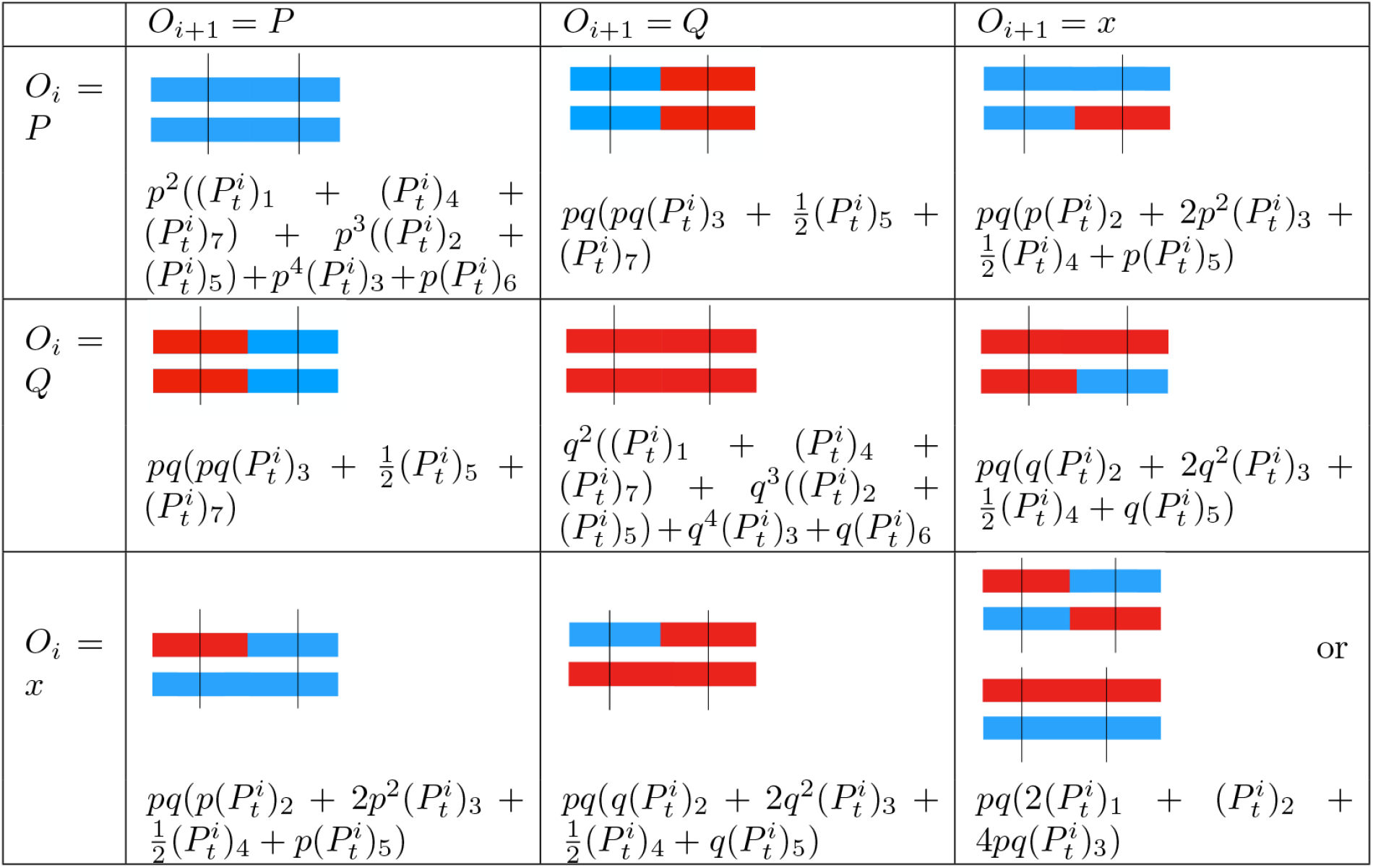
ℙ_*t*_(*O*_*i*_, *O*_*i*+1_) for unphased data. The allele from source taxa 𝒫 is represented in blue and the allele from source taxa 𝒬 is represented in red.

### 2.4 Individual based simulations

To test the validity of our maximum likelihood approach, we use individual based simulations, as described in (Janzen et al., 2018), i.e. Wright-Fisher type simulations of randomly mating populations of constant size *N*, with non-overlapping generations. We then recover local ancestry by analyzing ancestry at *n* markers whose positions are chosen uniformly at random along the genome.

Unless otherwise specified, we show how time can be accurately inferred for a population of 10, 000 individuals, for time points between the first generation and 1,000 generations. We use *n* = 10, 000 markers, which is considered to be sufficient to detect the majority of accumulated junctions (Janzen et al., 2018). We report our findings across 100 replicates, where in each replicate 10 individuals were randomly selected from the admixed population and used to infer *T*. We have simulated with three different values of the initial proportion of source taxa 𝒫, (*p* ∈ {0.053, 0.184, 0.5*}*), to vary the initial heterozygosity *H*_0_ in *{*0.1, 0.3, 0.5}.

We have first compared the methods we have developed here to previous methods based on the theory of junctions (Figure 5). We observe that, when the number of markers is low, previous methods, that do not take into account marker spacing, tend to underestimate the time since admixture, which is not the case for our methods.

**Fig 5.**
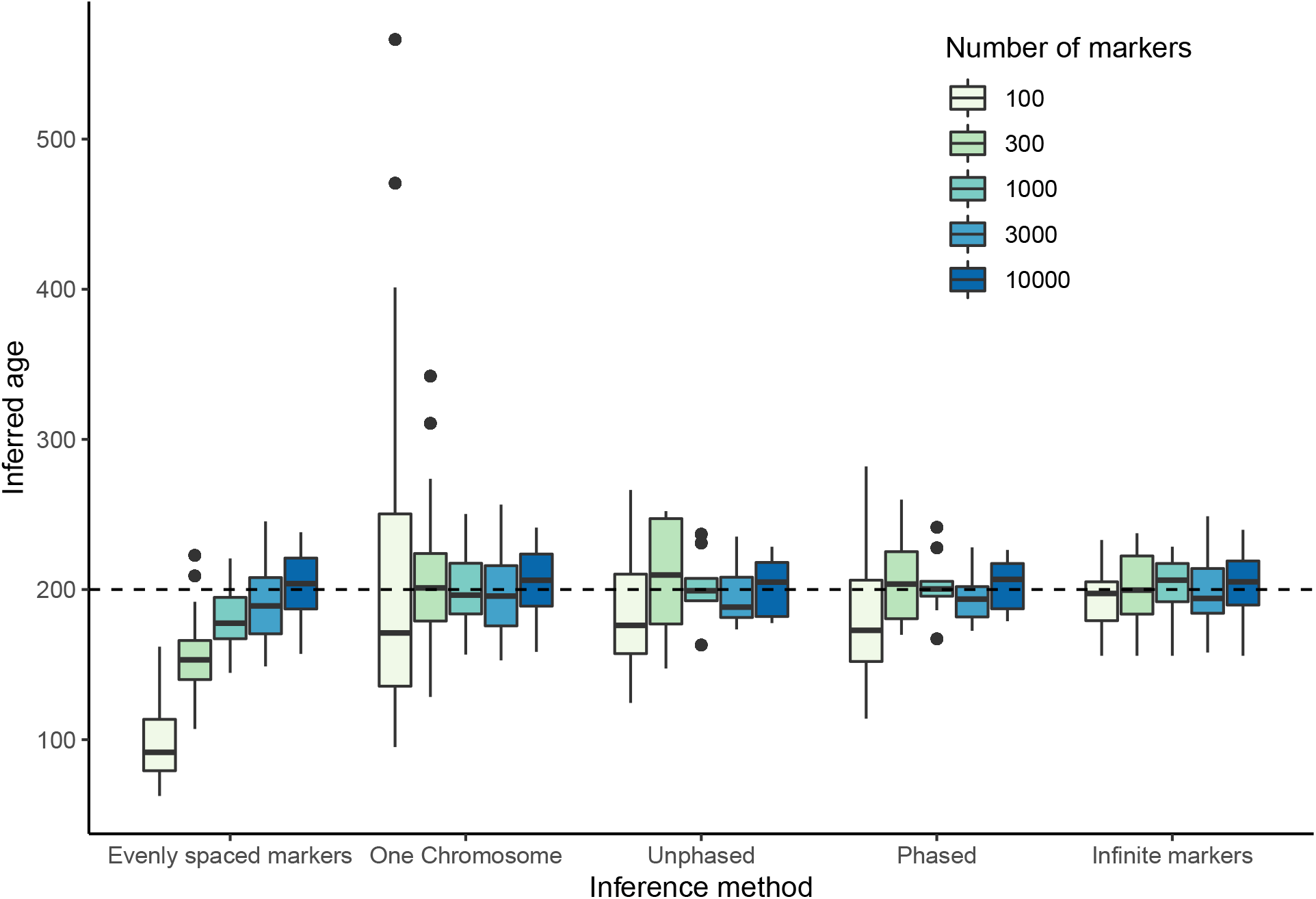
Comparison to previous methods. Shown are the median estimates for the time since admixture (dots) for 100 replicates, where in each replicate 10 individuals were analyzed. The dashed line indicates the simulated time. ‘Evenly spaced markers’ corresponds to the method in (Janzen et al., 2018). ‘Infinite markers’ corresponds to an idealized scenario where ancestry is known for every locus in the chromosome and is there to quantify the amount of randomness in the process. The population size was 10,000 individuals, and 10,000 randomly spaced markers were used.

We have then compared the estimations of the time since admixture, using the method for one chromosome and the method for two chromosomes (phased) (Fig 6). We observe that using data from the two homologous chromosomes allows to infer the time since admixture more accurately, since it reduces uncertainty.

**Fig 6.**
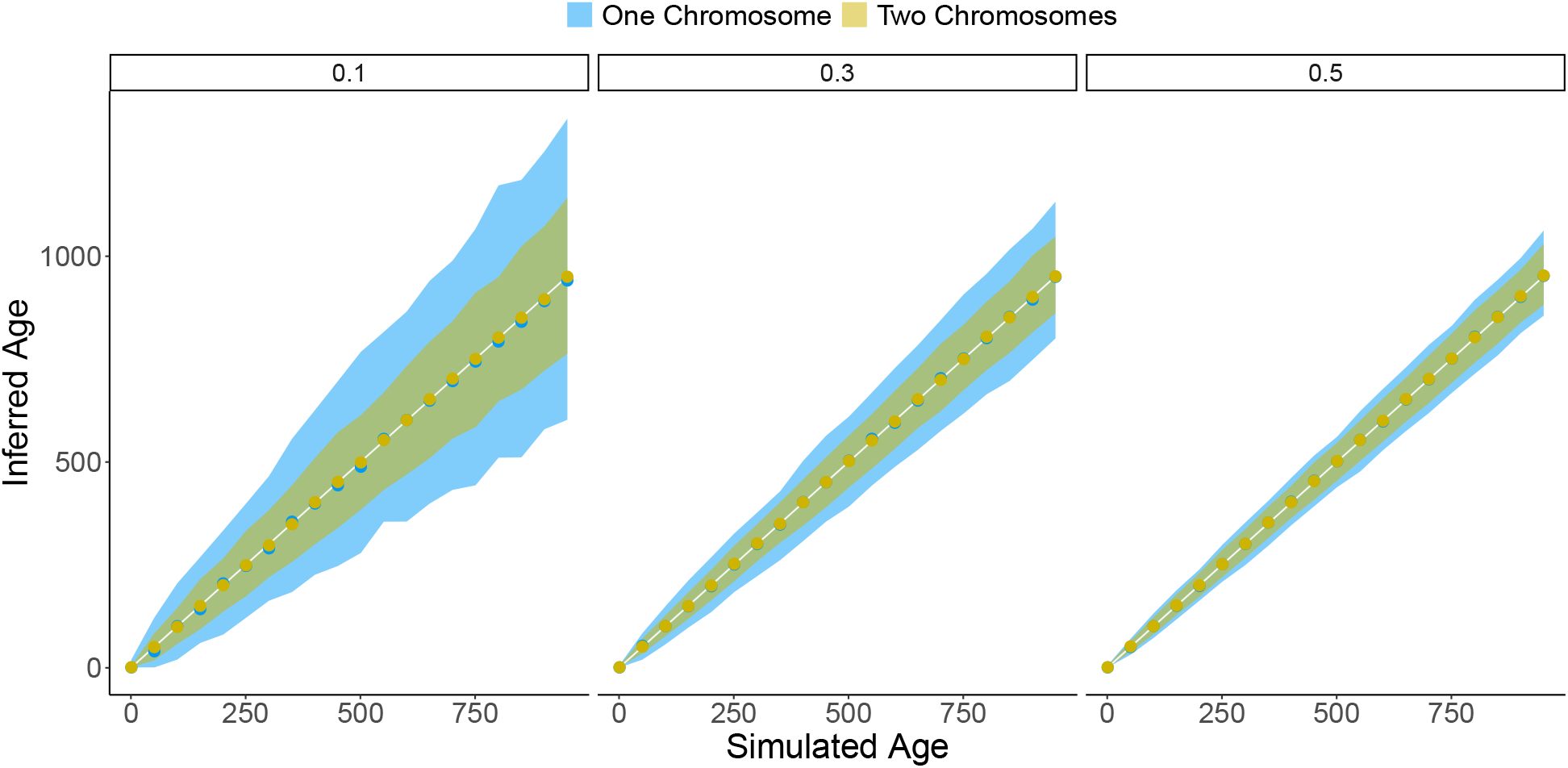
Accuracy in age estimate using information from one versus two chromosomes. Inferred time versus simulated time is represented. Shown are the median estimates (dots) for 100 replicates, where in each replicate 10 individuals were analyzed. The solid white line indicates the observed is equal to expected line and the shaded area indicates the 95% percentile. Shown are results using junction information from one chromosome (blue) and results using phased information from two chromosomes (gold). Numbers above the plots indicate the initial heterozygosity. The population size was 10,000 individuals, and 10,000 randomly spaced markers were used.

Using unphased data instead of phased data might introduce additional error and in Fig 7 we compare the methods that use phased or unphased information of two homologous chromosomes. We observe that both methods yield very similar results in terms of the relative error. This can be due to the fact that homozygous sites have an important contribution to the likelihood and the uncertainty that comes from sites that are of type *x* (in the unphased case) is well managed by our method.

**Fig 7.**
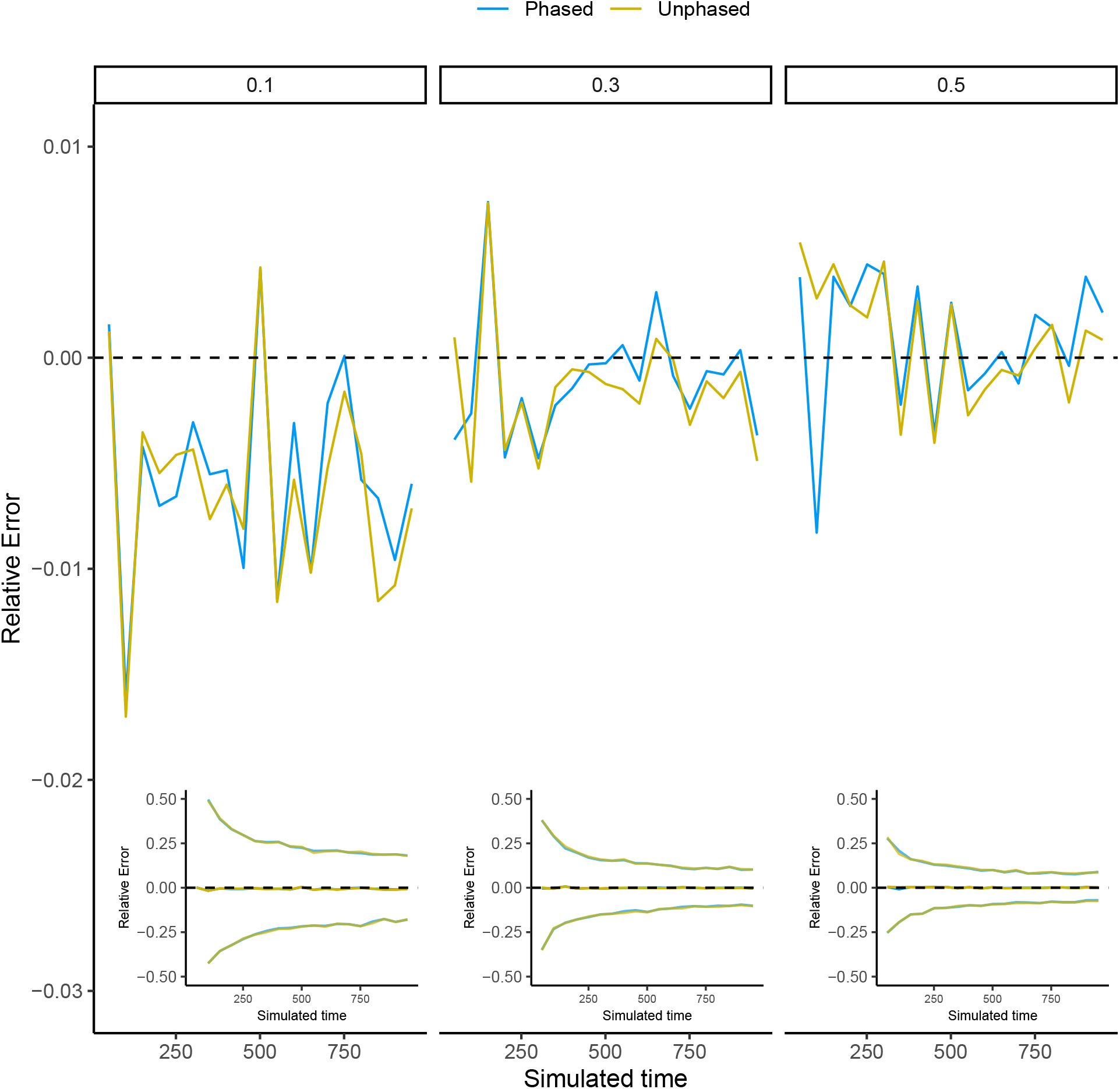
Accuracy in age estimate using the unphased framework versus the phased framework. Shown are the median difference across 100 replicates. We represent the results for three different initial heterozygozities, as indicated at the top of each plot. The population size was 10,000 individuals, and 10,000 randomly spaced markers were used. The inset plots show the same results, including the 95% confidence limits, which are far outside the boundaries of the main plot.

Finally, we explore error in phasing assignment (switching error). We simulate the effect of error in phasing assignment by randomly swapping a fraction of the markers between chromosomes. We explore phasing error in *{*0.0025, 0.005, 0.0075, 0.01, 0.02*}*. These errors are comparable to the switching error rates reported in the literature. For example, Choi et al. (2018) compared different phasing methods and reported switching error rates between 0.1% and 2%. (Notice that these error rates are for human data where there are good quality references and sample sizes are large). More recent reference-free methods (based on third generation sequencing techniques) report switching error rates of 1-2% (see for example Tourdot and Zhang (2019); Ebler et al. (2019); Kronenberg et al. (2019)). Switching error rate error has strong effects on the inferred time since admixture, as shown in the bottom panel in Fig 8. Generally, imposed errors increase the inferred age, by introducing novel junctions due to mis-phased markers.

**Fig 8.**
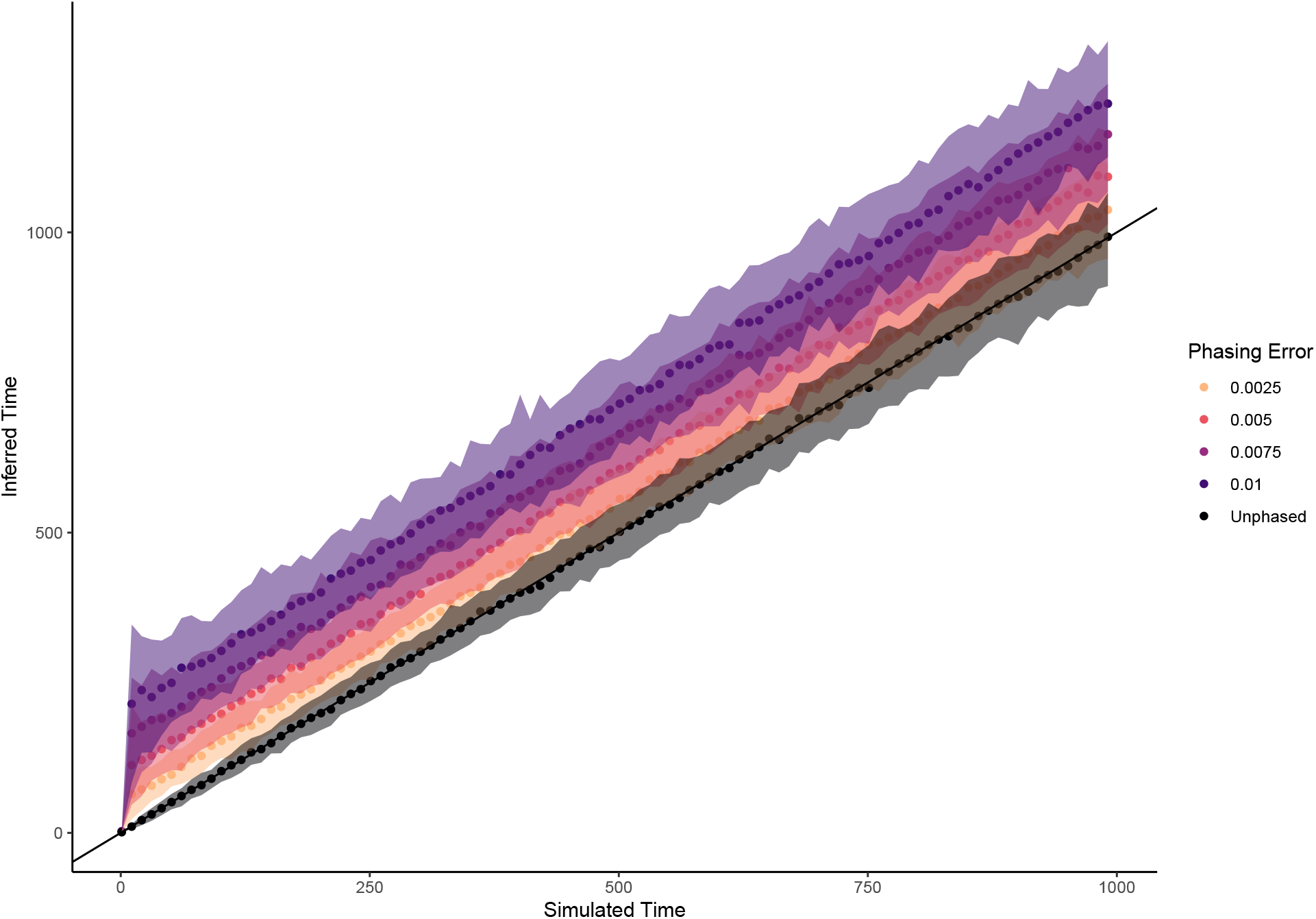
Effect of switching error on the estimated time since admixture. Data simulated with *N* = 10, 000, *p* = 0.5, *C* = 1 and *n* = 10, 000. The solid black line indicates the simulated = estimated time. Dots indicate the median inferred age and the colored area indicates the 95% confidence interval (CI) envelope. Colors reflect different degrees of phasing error, where a phasing error of 0.01 represents a 1% probability of a SNP being phased incorrectly.

Another important source of error is the lack of coverage, which would have the effect of reducing the number of markers. An analysis of the sensitivity of our method to reducing the number of markers can be found in the S3 Appendix, and shows that a reduction of coverage also tends to lead to an overestimation of the time since admixture.

## 3 Comparison with other methods

Our framework provides a methodology to infer the time since admixture, provided that local ancestry is known. Previous methods aiding in inferring the time since admixture, such as ELAI (Guan, 2014) and ancestry hmm (Corbett-Detig and Nielsen, 2017) jointly infer the time since admixture and local ancestry, because they use the time since admixture to correct for the impact of recombination on local ancestry estimates. We have chosen to compare our framework with results obtained using ancestry hmm, which we currently consider the most accurate method available to jointly infer local ancestry and time since admixture. We do so in a two-fold manner: first, we directly compare the ability of both methods to infer the time since admixture, provided data with known local ancestry. Secondly, we provide both packages with imperfect data, where local ancestry is uncertain. Because our framework is not able to infer local ancestry, we directly use the inferred local ancestry of ancestry hmm, which might add an extra layer of error, but is a good reflection of an empirical use-case (see also section 4.3). For both approaches (known and uncertain ancestry), data was simulated using the junctions package, using a population size *N* of [1000, 10000] and imposing *n* markers, where *n* was [1000, 10000, 40000] (in line with the number of markers available in the Swordtail and *Populus* datasets in section 4). Simulations were run for 10,000 generations, using *H*_0_ = 0.5 and *C* = 1. At each generation, time since admixture was inferred for 10 individuals sampled randomly from the population. Time since admixture was inferred given the superimposed *n* markers, whose location was drawn randomly in [0, 1]. For each parameter combination, 10 replicate simulations were performed. When the maximum likelihood did not converge to a final value within the given range, the data point was removed from the analysis.

### 3.1 Known ancestry

Because ancestry hmm requires information on allele frequencies in the source taxa in order to jointly infer local ancestry and time since admixture, input data for ancestry hmm was created assuming both source taxa were fully separated, e.g. two alleles were differentially fixed across the source taxa. Results show (Figure 9) that in the limit of large *N*, our method performs as good as ancestry hmm. However, for smaller population sizes, we observe that ancestry hmm becomes increasingly incorrect as the time since admixture increases, in contrast to our method, which remains accurate.

**Fig 9.**
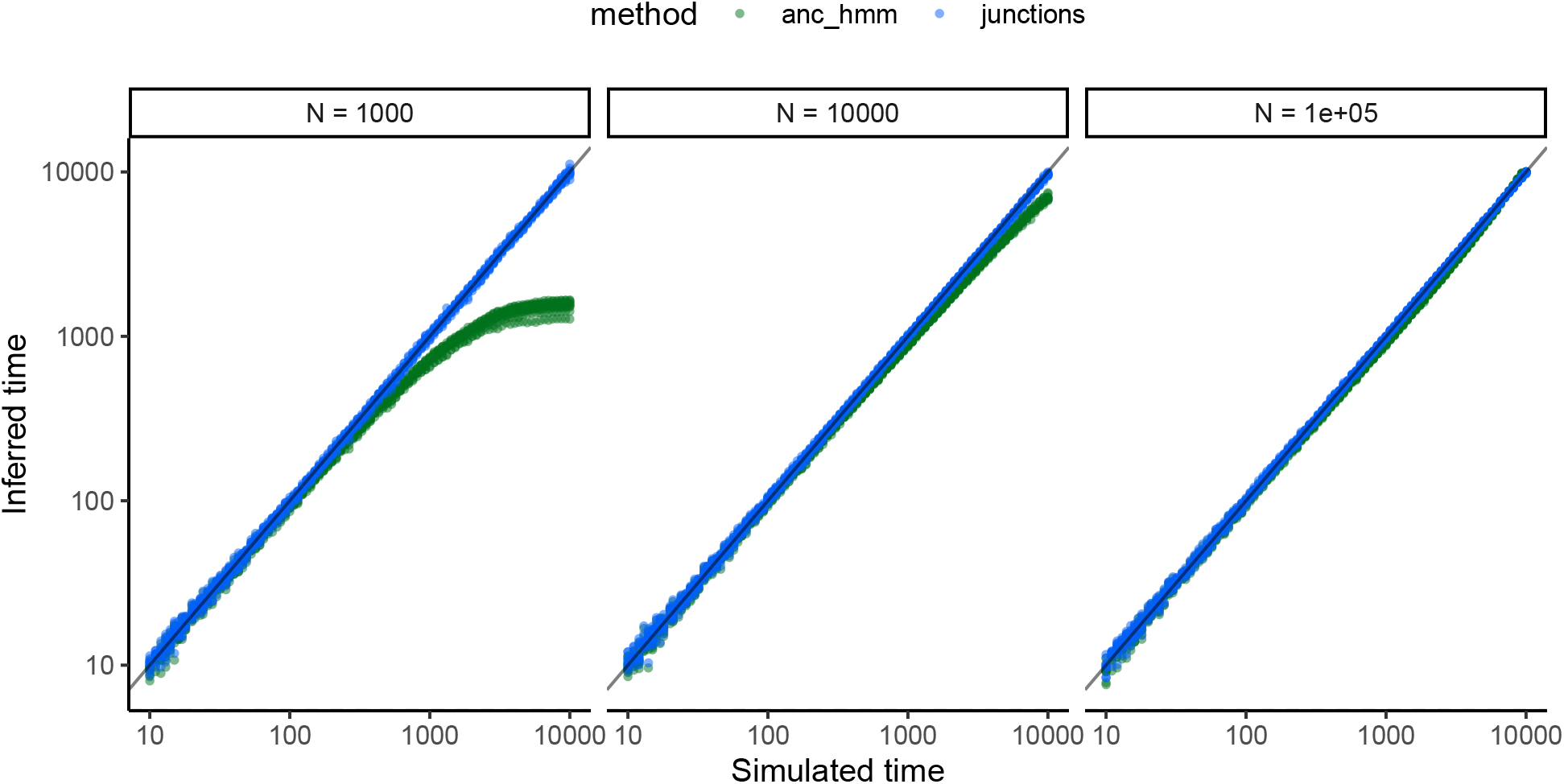
Comparison in estimating the time since admixture between ancestry hmm and the method proposed here. The solid black line indicates the simulated = estimated time. Dots indicate the inferred ages, with the green dots representing ages inferred by ancestry hmm and the blue dots indicate ages inferred by the junctions framework. Age estimates are based on simulated data with known ancestry, using *n* = 40, 000, *C* = 1, *H*_0_ = 0.5.

### 3.2 Uncertain ancestry

To mimic uncertainty in ancestry, we explored different degrees of uncertainty, indicated by different values of the allele frequency differential *d*_*x*_ = *A*_*x*_ − *A*_*y*_, where *A*_*x*_ is the frequency of allele *A* in population *x* and *A*_*y*_ is the frequency of allele *A* is population *y*, following Shriver et al. (1997). Higher values of *d*_*x*_ indicate higher ancestry information content in the respective marker. Here, we explored values of *d*_*x*_ in [0.5, 0.7, 0.9]. Because our framework does not infer local ancestry, we used the inferred local ancestry by ancestry hmm, using a threshold of including only markers with at least 95 % certainty of ancestry. For a large population size (*N* = 10, 000, Supplementary material S5) we observe that ancestry hmm consistently underestimates the true age when the time since admixture becomes large, in line with our findings with known ancestry (Figure 9). Our method underestimates the true age even further, most likely as a result of a stacking of errors: because our method can not infer local ancestry, we have used the local ancestry estimates of ancestry hmm. However, when population size is small (*N* = 1000, Figure 10), we observe a striking phenomenon (Figure 10): when the time since admixture becomes large, our framework starts inferring the time correctly again, whereas the age estimates obtained using ancestry hmm remain incorrect and seem to reach a plateau. This effect only occurs when there is a sufficient number of markers of reasonable quality, as we observe that when *n* = 1000 and *d* ≤ 0.7, our framework is also unable to accurately infer the time since admixture. We believe that the improved performance when time since admixture becomes large is the result of the interplay between two phenomena. Firstly, while our method can work for any value of *T*, ancestry hmm typically requires *T* ≪ log(*N*) (this is the same assumption used in SMC’ (Liang and Nielsen, 2014)). This could introduce some error in the ancestry assignment and thus lead to the underestimates obtained using ancestry hmm. Secondly, our method seems to work better with a small number of informative markers. Combined, our method outperforms ancestry hmm in inferring the time since admixture when population size and number of markers is small.

**Fig 10.**
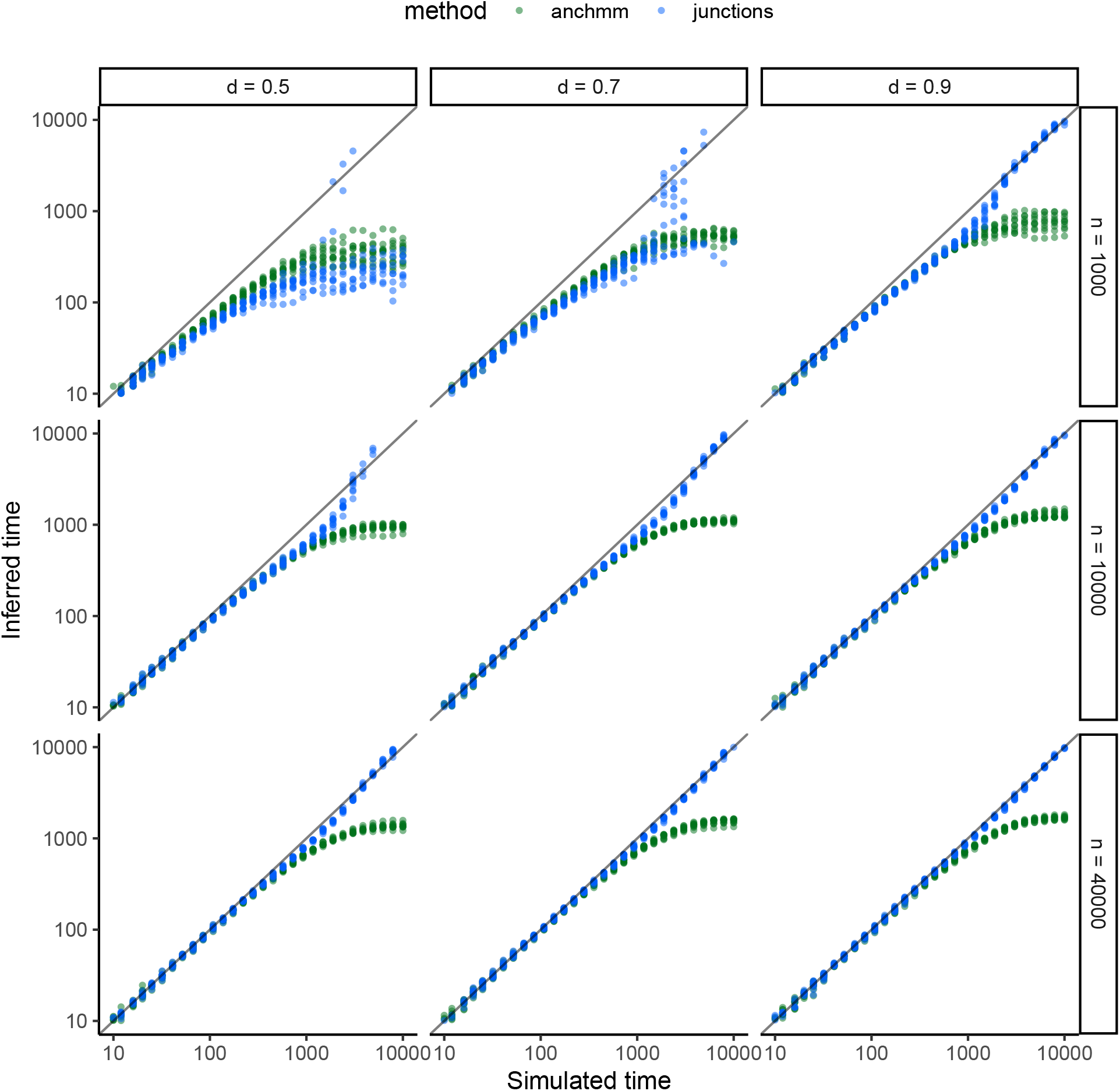
Comparison in estimating the time since admixture between ancestry hmm and the method proposed here, for a small population of *N* = 1000.. The solid black line indicates the simulated = estimated time. Dots indicate the inferred ages, with the green dots representing ages inferred by ancestry hmm and the blue dots indicate ages inferred by the junctions framework. Missing dots are caused by a lack of convergence of the maximum likelihood algorithm. Age estimates are based on simulated data with uncertain ancestry, where uncertainty in ancestry is reflected by the allele frequency differential (Shriver et al., 1997). Because the method proposed here does not include ancestry uncertainy, local ancestry as inferred by ancestry hmm was used.

## 4 Results

### 4.1 Saccharomyces cerevisiae

Experimental evolution provides an important reference point to verify our findings. Here, we reanalyze data from an Advanced Intercross Line (AIL) experiment, where two highly differentiated yeast (*Saccharomyces cerevisiae*) lines were crossed, and the resulting hybrid offspring was outbred for 12 generations in order to obtain maximum genetic diversity (Parts et al., 2011; Illingworth et al., 2013). The data consists of sequencing data for 171 individuals, for all 16 chromosomes. There are on average 3271 ancestry informative markers per chromosome (95% CI: [929, 6284]). Local ancestry was certain in the data, as both source taxa were differentially homozygous on each ancestry informative SNP. *H*_0_ was 0.5, reflecting a 50/50 contribution of both strains to the first generation. We used three different recombination rate estimates: firstly, we used the linkage map of Cherry et al. (1997) where the average recombination rate is 1*cM/*2.7*kb* (1 centimorgan per 2.7 kilobases), secondly, we used the average recombination rate of 1*cM/*2.2*kb* as inferred in Mancera et al. (2008) and lastly, we used the average recombination rate of 1*cM/*5.8*kb* as inferred for the two-way cross in Illingworth et al. (2013). In the absence of a detailed recombination map, we assume that recombination is constant across the chromosome, ignoring hotspots and coldspots. We assume a large population size (*N* = 100, 000), reflecting outbreeding.

We find that when using the older recombination rate estimates, we consistently underestimate the age of the hybrids (See Figure 11, panel A). The mean age using the recombination rate from Cherry et al. (1997) corresponds to 9.1 generations (95 % CI across all individuals = [6.2, 11.3]) and the mean age using the recombination rate from Mancera et al. (2008) corresponding to 7.0 generations (95 % CI: [5.4, 8.6]). These estimates suggest that the true recombination rate is slightly lower than assumed. When using the most recent recombination rate estimate (i.e. 1*cM/*5.8*kb*, from Illingworth et al. (2013)), we slightly overestimate the age (mean age estimate: 18.7 generations, 95 % CI: [13.0, 22.5]). Alternatively, we could be overestimating population size, suggesting that perhaps the rate of inbreeding in the experimental design was higher than anticipated. Reducing the population size tends to increase the time since admixture (Supplement S4), which could help bring the age estimates using the two oldest recombination rates closer to the true number of generations. However, for the most recent recombination rate estimate, this would only increase the error made. The likelihood profiles (See Figure 11, panel B) suggest that the maximum likelihood estimates are robust and thus it seems more likely that the recombination rate estimates do not accurately reflect the amount of recombination accumulated in the experiment.

**Fig 11.**
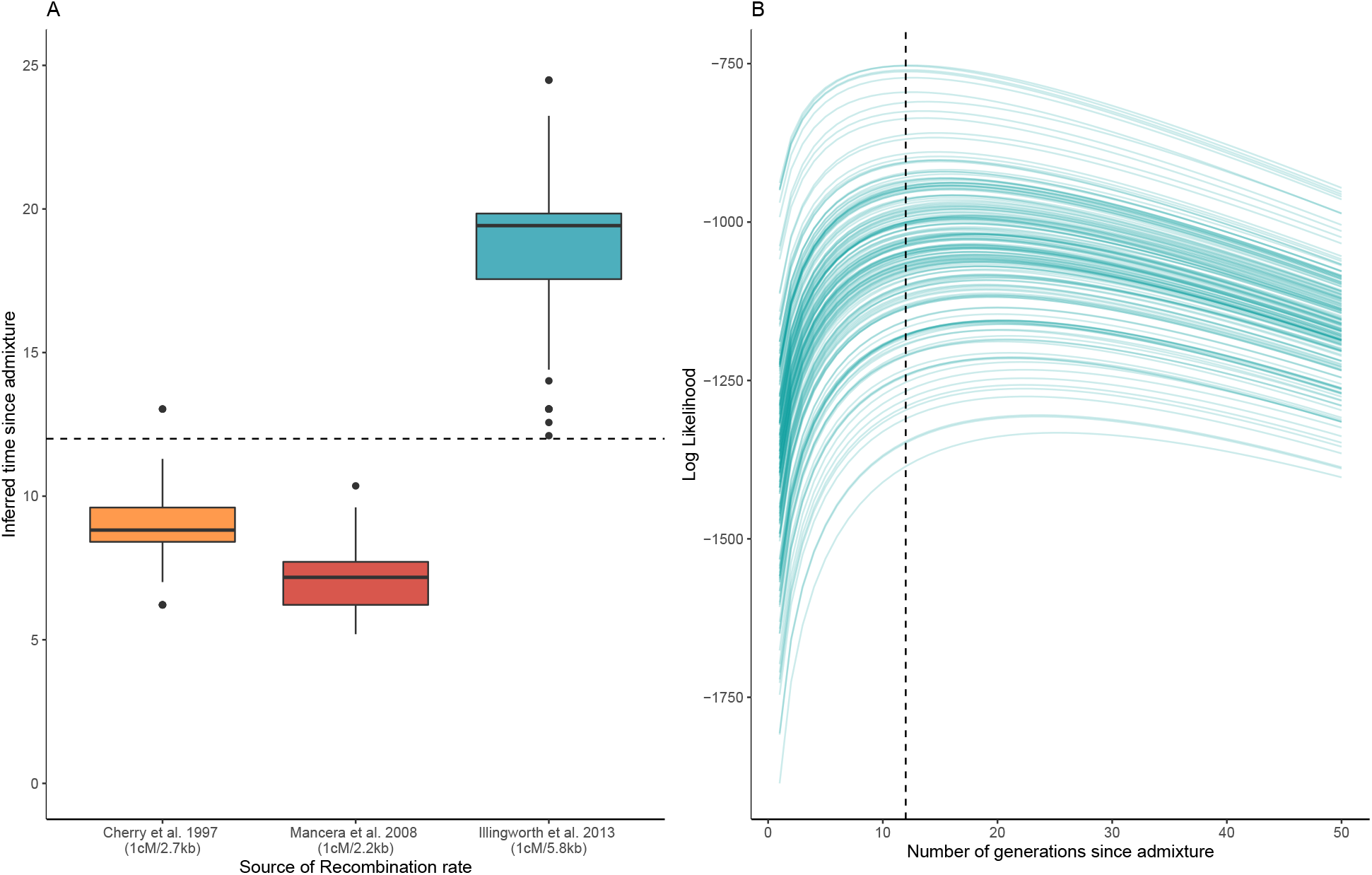
Inferred age for F12 Hybrid Yeast (*Saccharomyces cerevisiae*) individuals. (A) Inferred age for three different recombination rates: 1*cM/*2.7*kb* (Cherry et al., 1997), 1*cM/*2.2*lb* (Mancera et al., 2008) and 1*cM/*5.8*kb* (Illingworth et al., 2013). Shown is the distribution of inferred ages across 171 individuals. (B) Loglikelihood profile across all chromosomes, where each line represents one individual. Shown is the loglikelihood profile for the most recent recombination rate estimate, from Illingworth et al. (2013). The vertical dotted line indicates the 12 generations line (e.g. the true age of the F12 Hybrid individuals).

### 4.2 Swordtail Fish

Here, we reanalyze data of hybridizing swordtail fish published in (Schumer et al., 2018). Swordtail fish have received considerable attention in the past years, as they have been shown to hybridize readily in nature. We focus here on a hybrid population located in Tlatemaco, Mexico (Schumer et al., 2018, 2014a). The population is the result of a hybridization event between *Xiphophorus birchmanni* and *X. malinche*, approximately 100-200 generations ago (Pers. Comm. M. Schumer and (Schumer et al., 2018; Powell et al., 2020a)). Currently, the hybrid genome consists for 75% of *X. malinche*, suggesting that the initial hybrid swarm was strongly biased towards *X. malinche*, or that strong selection after hybridization has favored genomic material from *X. malinche*. We use ancestry information provided in the data supplement of Schumer et al. (2018), which contains unphased local ancestry estimates based on multiplexed shotgun genotyping (MSG) results (Andolfatto et al., 2011), with on average 38,936 markers per chromosome (95% CI: [22765, 50256]). The MSG pipeline provides a posterior probability of observing local ancestry. Following Schumer et al. (2018), we converted local ancestry probabilities of *>*95% to hard ancestry calls. To obtain age estimates, we use the estimated population size in Schumer et al. (2014a) of 1830 individuals. We infer the age for 187 individuals from the Tlatemaco population. We use ancestry information from 24 linkage groups, but remove linkage groups 17 and 24, as these are known to include large inversions (Schumer et al., 2018), making them unsuitable for admixture analysis. As a recombination map, we use three approaches. Firstly, we use the average recombination rate of 1*cM/*378*kb* as used in Schumer et al. (2014a), which is based on the average genome-wide recombination rate in *Xiphophorus* (Walter et al., 2004). Secondly, we use the average recombination rate of 1*cM/*500*kb* as reported in Powell et al. (2020b). Lastly, we use the high density recombination map reconstructed from Linkage Disequilibrium patterns as presented in Schumer et al. (2018), which represents an average recombination rate of 1*cM/*485*kb*.

We find that the distribution of ages inferred for individuals from the Tlatemaco population is overall higher than the previously inferred age but still consistent with those estimates (see Fig 12 A). We recover a mean age of 163 generations (95% CI: [81, 201]) when using the recombination rate reported in Schumer et al. (2014b). Using the high density recombination map from Schumer et al. (2018) we obtain a mean age estimate of 210 generations (95% CI: [111, 257]), due to the shorter map length. Alternatively, using the most recent recombination rate estimate of 1*cM/*500*kb* reported by Powell et al. (2020b), we recover a mean age of 217 generations (95% CI: [115, 265]). Thus, we generally find that the population is perhaps slightly older than expected, but we would like to emphasize that the likelihood profile across all samples (Figure 12 B) is extremely flat, suggesting that the age estimates obtained tend to be uncertain and susceptible to potential inconsistencies in the data (e.g. sequence errors). Exploration of different values for the population size (Supplement S4) shows that if the true population size is larger than reported by Schumer et al. (2014a), this would decrease the time since admixture, but only minimally so (by only a few generations).

**Fig 12.**
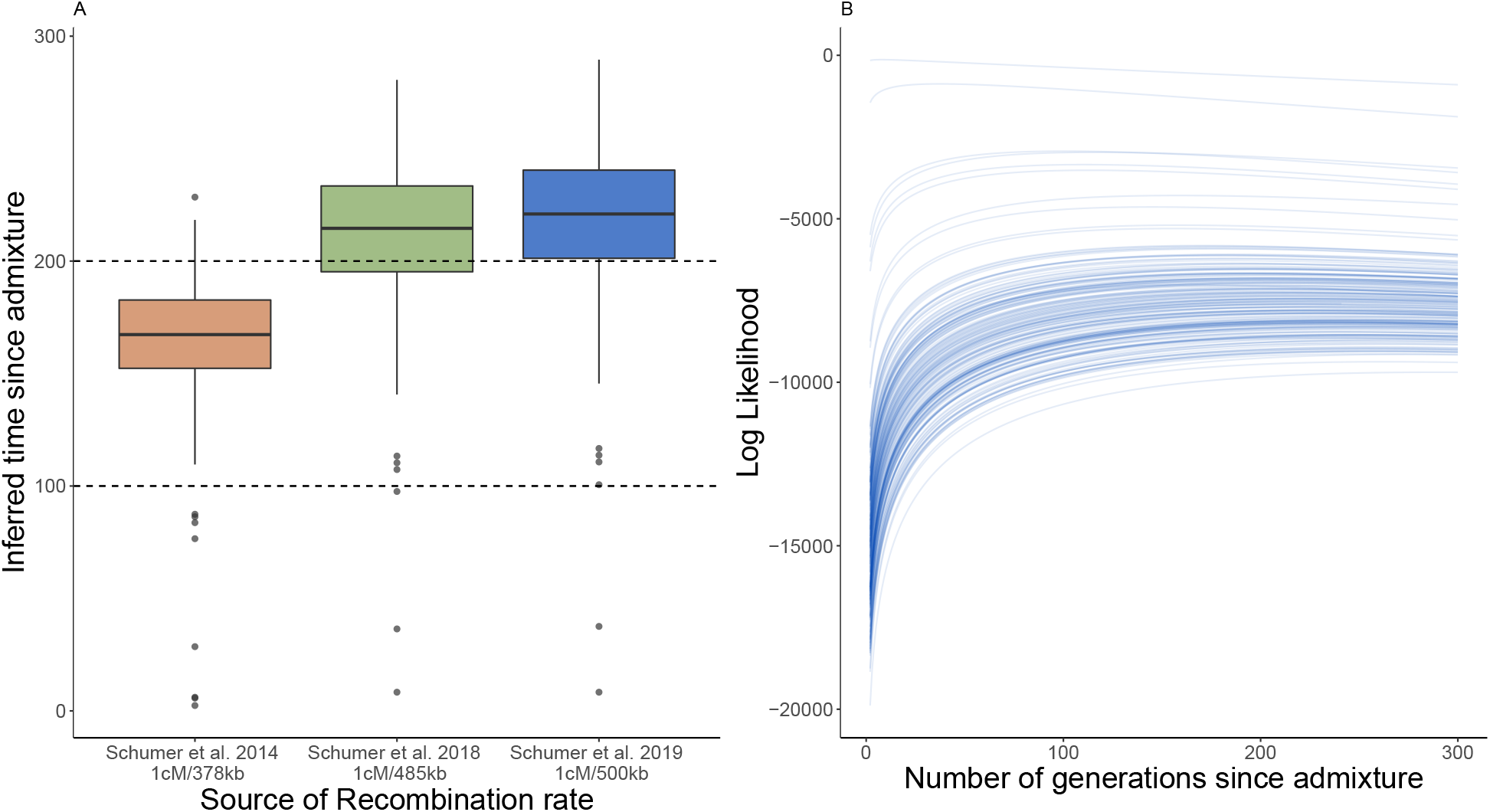
Inferred age for hybrid *Swordtail* fish. (A) Inferred age for hybrid *Xiphophorus* fish from Tlatemaco (Mexico). Shown are age inferences based on three different recombination maps: 1*cM/*370*kb* (Schumer et al., 2014a), 1*cM/*485*kb* (Schumer et al., 2018) and 1*cM/*500*kb* (Powell et al., 2020b). Dotted lines indicate the hypothesized age limits of the admixed population. (B) Loglikelihood profile across all chromosomes, where each line represents one individual. Shown is the loglikelihood profile for the most recent recombination rate estimate (Powell et al., 2020b).

### 4.3 *Populus* trees

Here, we reanalyze a dataset of *Populus* trees, published by Suarez-Gonzalez et al. (2016). The dataset focuses on two species of trees, *P. trichocarpa*, found mainly in West-America, and *P. balsamifera*, which is found in Northern America. The two species are thought to have diverged relatively recently, around 760k years ago. Where their ranges meet (around the southern tip of Alaska), the two species hybridize, and a hybrid population has been established. The dataset consists of 32 individuals which are mainly *P. balsamifera*, admixed with *P. trichocarpa* and 36 individuals that are mainly *P. trichocarpa*, admixed with *P. balsamifera*. Three chromosomes of interest (chromosomes 6, 12 and 15) were Illumina sequenced, and unphased data was available for on average 60071 ancestry informative markers per chromosome (95% CI: [28745, 101425]). Ancestry information in the markers was very high, with on average 40852 markers with an allele frequency differential of at least 0.7 (Shriver et al., 1997), and 21009 markers with an allele frequency differential of at least 0.9. We use three different population level recombination rates recovered from the literature, being *ρ* = 0.00219 (Wang et al., 2016), *ρ* = 0.0092 (Olson et al., 2010) and *ρ* = 0.0197 (Slavov et al., 2012). We converted these population level recombination rates to individual rates using an effective population size of 5106 individuals, as estimated using phylogenetic methods in (Slavov et al., 2012). This yielded three local recombination rates of 1*cM/*10.4*kb* (Slavov et al., *2012), 1cM/*22.2*kb* (Olson et al., 2010) *and 1cM/*93.3*kb* (Wang et al., 2016). Local ancestry was determined using ancestry hmm (Corbett-Detig and Nielsen, 2017), assuming equal admixture of both source taxa. Because admixture differed strongly across samples, we used the average local ancestry per sample as input for a second run of ancestry hmm in order to obtain accurate local ancestry calls. We converted local ancestry probabilities of >95% to hard ancestry calls. Lastly, we compared the observed variation in ancestry across samples with the expected variation in ancestry for a single panmictic population, as given by equation 8 in (Gravel, 2012).

We find that the time since admixture strongly correlates with the recombination rate used (See Figure 13 A), with a mean number of generations since admixture of 6 (95% CI: [3, 13]) when using the highest estimate of recombination (1*cM/*10.4*kb* (Slavov et al., 2012)), an intermediate estimate of 12 generations (95% CI: [7, 28]) when using a recombination rate of 1*cM/*22.2*kb* (Olson et al., 2010) and a much higher age estimate of 50 generations (95% CI: [30, 114]) when using the lowest recombination estimate of 1*cM/*93.3*kb* (Wang et al., 2016). The likelihood profiles (See Figure 13 B) intersect on multiple occasions, suggesting multiple optima. We measure a variation in ancestry across the three chromosomes of 0.042. However, we obtain expected levels of variation of ancestry of 0.0088, 0.0010 and 0.0008 for the three different recombinatinon rate estimates (and their corresponding age estimates). Observed variation in the data is thus much higher than expected from admixture in a single panmictic population.

**Fig 13.**
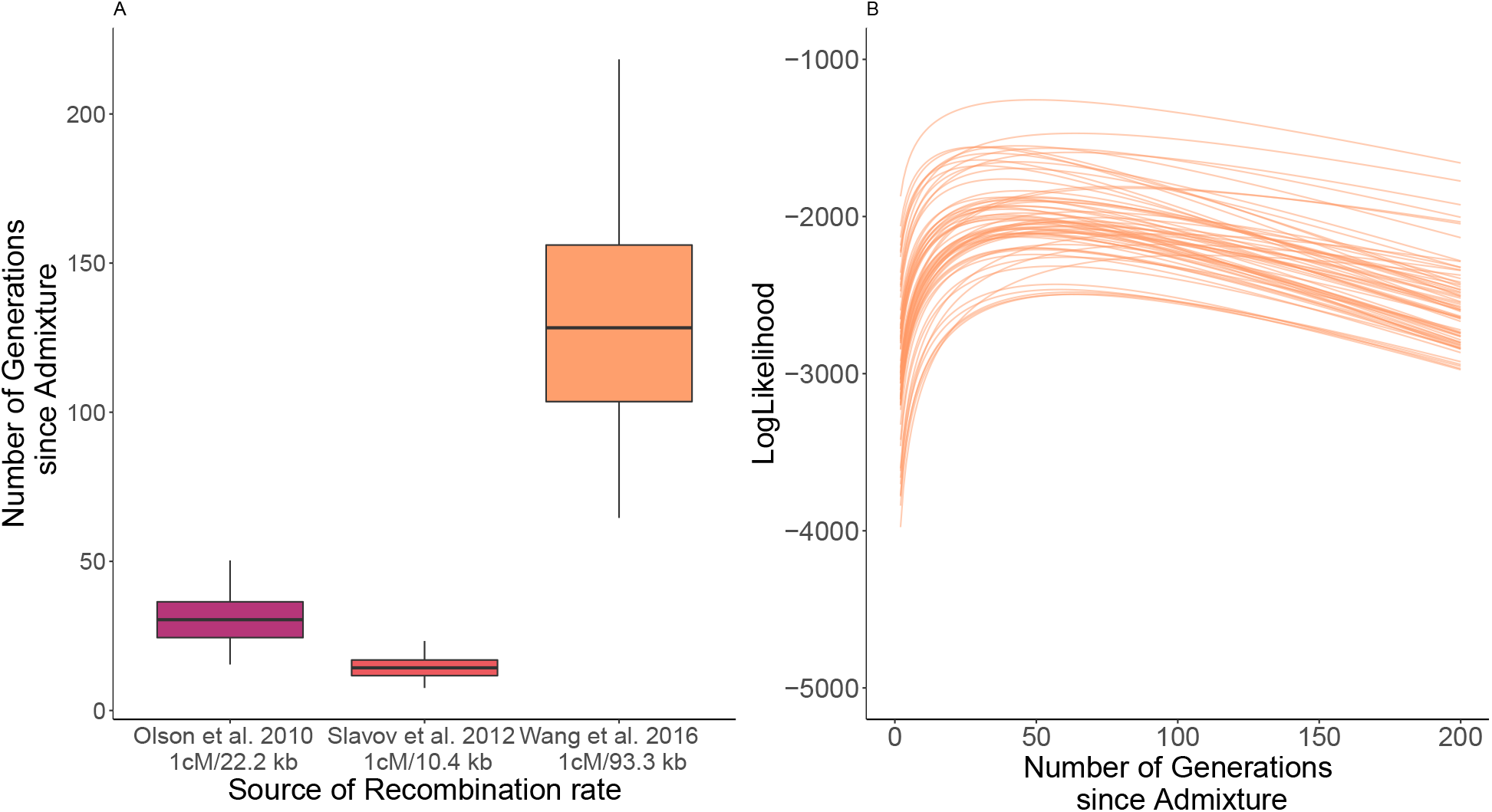
Inferred age for hybrid *Populus* trees. (A) Distribution of inferred time since admixture for all individuals. Colors indicate different recombination rates used: 1*cM/*10.4*kb* (Slavov et al., 2012), 1*cM/*22.2*kb* (Olson et al., 2010) and 1*cM/*93.3*kb* (Wang et al., 2016). (B) Inferred time since admixture, split out across the average frequency *of P. trichocarpa* in the admixed individual. (C) Loglikelihood profile across all chromosomes, where each line represents one individual. Shown is the loglikelihood profile for the most recent recombination rate drestimate (Wang et al., 2016).

## 5 Discussion

The aim of this article was to improve the estimation of the time since admixture in hybrid populations. To do so, we have extended the theory of junctions in two directions. First, we have derived a formula for the expected number of observed junctions in one chromosome that takes into account the number of markers and their positions (equation (3)). Second, we have considered the case in which there is sequencing data from two homologous chromosomes. We have developed a maximum likelihood approach that infers the time since admixture, for both phased and unphased data.

Firstly, we have used simulations to validate the accuracy of our method. Results from our simulations show that our method for a single chromosome performs better than previous methods that ignore the effect of having a limited number of markers or assume that the markers are even-spaced (see Fig 5). Furthermore, we find that using information from two chromosomes improves accuracy considerably, as expected. This effect is even stronger for small population sizes (see Fig 6 and Fig 2 in S2 Appendix).

Surprisingly, the method based on unphased data performs similarly to our method based on phased data (see Fig 7 and Fig 3 in S2 Appendix). The phased and unphased approaches differ in their treatment of markers that are heterozygous for ancestry, and hence we expected differences between these methods to manifest themselves primarily during the initial stages of admixture, when heterozygosity is still high. We did find that there were slight differences during these stages (Fig 7 and Fig 3 in S2 Appendix), but these were neglibile compared to the overall uncertainty. Furthermore, these errors were much smaller than errors accumulated due to incorrect phasing (see Fig 8). Our findings here are conservative, as we show that the unphased method performs better even for small error rates, comparable to error rates for human data (for example in (Choi et al., 2018)). Human data sets are typically of very high quality, and these error rates represent an extremely favourable scenario. Thus, given the impact of error rates incurred during phasing, we recommend using our unphased framework if possible, to obtain more accurate time estimates.

Apart from sensitivity to phasing error, we have tested the sensitivity of our method to different parameters such as the number of markers *n*, the population size *N*, the initial heterozygosity *H*_0_ and the total recombination rate *C* (see S1 Appendix). Our method seems to be quite sensitive to *H*_0_ but this parameter can easily be estimated from the proportion of markers that come from each source taxa. One advantage of our approach is that age inference is not very sensitive to population size (see Fig 1 in S1 Appendix), which was not true for previous methods that rely on a good estimation of *N* (see Janzen et al. (2018)). Our method is not very sensitive either to the number of markers (see Fig 4 in S1 Appendix), provided that it is above a certain threshold. Janzen et al. (2018) inferred that when using regularly spaced markers and information for a single chromosome, the number of markers typically needs to be an order of magnitude larger than 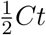, where *t* is the admixture time and *C* the total amount of recombination. We find similar results when using information from a single chromosome with arbitrarily spaced markers or information from both chromosomes (see S1 Appendix). When analyzing empirical data, it is often impossible to know *a priori* whether the number of ancestry informative markers is much larger than the admixture time. However, our simulation results indicate that when the number of markers is too small, variation in the age estimate across different chromosomes tends to increase. Thus, large variation in the estimate of admixture time, or inferred admixture times that tend to extremely large values, are potentially indicative of an insufficient marker number.

The main issue with our method is its sensitivity to the recombination rate. This is shown in Fig 2 of S1 Appendix but also exemplified by the varying results in the empirical datasets, dependent on our assumptions about recombination rates. However, it should be noted that this issue is not novel to our approach, but is a general issue with the theory of junctions. Apart from sensitivity to the average recombination rate, local hot-spots or cold-spots of recombination could potentially also influence admixture time estimates.

Furthermore, the recombination rate factors into inference of local ancestry. Our methodology assumes local ancestry to be known, and relies on upstream inference of local ancestry. However, methods to infer local ancestry such as MSG (Andolfatto et al., 2011), ELAI (Guan, 2014) and ancestry hmm (Corbett-Detig and Nielsen, 2017) also use recombination in their calculations to infer local ancestry. Thus, by using these local ancestry estimates, our methodology re-uses the recombination rate (both for local ancestry inference, and subsequent age estimation). Any errors in the recombination rate estimate might then propagate through the pipeline and affect age estimates using our framework. Using simulations including ancestry uncertainty, we have looked into this effect of stacking errors and have found that our method performs similarly to ancestry hmm, even if we use the local ancestry inferred by ancestry hmm. Furthermore, our method outperforms ancestry hmm for long admixture times or for small population sizes. ancestry hmm assumes *T* ≪ log_2_(*N*) (this is the same assumption used in SMC’ (Liang and Nielsen, 2014)), whilst our framework does not need to make these assumptions to obtain age estimates. However, when marker numbers dwindle (Supplement S5), our framework can no longer correct for this approximation, and performance of our framework drops below inference using only ancestry hmm. Yet, in the empirical datasets we have analyzed, we have generally found marker densities for which these limits were not reached and with full genome sequencing now within reach for many researchers, we expect this not to be an issue.

To validate our approach we have re-analysed three datasets. The first dataset is from a crossing experiment with *S. cerevisiae*. Here, we applied the single chromosome equations, and estimates of the time since the onset of admixture line up well with the experimental design, although assumptions regarding the recombination rate remain of strong influence on the admixture time estimates.

The second dataset we reanalyzed is of Swordtail fish (*Xiphophorus*). We infer an admixture time that is older than previous estimates (Schumer et al., 2014b) but that is in line with more recent estimates done by the same authors (M. Schumer, personal communication, (Powell et al., 2020a)) using more recent recombination rate estimates (Powell et al., 2020b). However, likelihood profiles are fairly flat, indicating that our maximum likelihood estimates are sensitive to changes in the data, further reflected by the sensitivity of the results to assumptions made regarding the recombination rate.

Finally, we have reanalyzed a dataset on *Populus* trees (Suarez-Gonzalez et al., 2016). We infer an admixture time that is in line with previous findings, but the original analysis did not focus on admixture time and only used admixture time to infer local ancestry. However, we find that the time since admixture correlates strongly with the genetic distance to either of the source taxa, with individuals more closely related to the source taxa inferred to be younger. This suggests that the dataset does not consist of a sample from a single, admixed, population and that this invalidates our analysis. Furthermore, variation in ancestry is much higher than expected from the admixture time alone (Gravel, 2012), again indicating that the analyzed population is most likely not a single admixing population. Lastly, admixture mapping analyses have shown that perhaps late generation backcrosses have contributed as well to the hybrid population (Suarez-Gonzalez et al., 2018), suggesting an intermediate form between on the one hand adaptive introgression & back-crossing and on the other hand ongoing hybridization across a spatial gradient. Across these results, it is clear that our assumption of a single admixed population is violated, and hence it seems likely that our age estimates are incorrect and that our framework is not a good fit for this empirical dataset. Yet, we believe that it provides an interesting example on how our methodology can be applied.

Summarizing, we have presented a framework to estimate the time since admixture using phased or unphased data from two homologous chromosomes, taking into account marker spacing along the chromosome. We have shown that using data from two chromosomes improves the estimations of the admixture time compared to the method that uses only one chromosome. This is true whether the data is phased or unphased. In addition we have shown, using simulations, that applying the phased or the unphased method yields very similar results. However, given that even small (unavoidable) phasing errors produce overestimates in the time since admixture, we suggest that, in most cases, using unphased data is the best strategy. In comparison with previous methods, such as ancestry hmm (Corbett-Detig and Nielsen, 2017), our method performs better when the population size is small or the time since admixture is long. With our new framework, we hope to have opened new avenues towards inferring the time since admixture in admixed populations, and primarily hope to have brought this analysis within reach also for datasets where phased data is unavailable or impossible to acquire. Furthermore, we would like to emphasize that our method also works for a relatively small number of SNPs, which opens up avenues towards analysis of closely related taxa, where the number of ancestry informative markers might be low. We have included the derivations and the numerical solution framework in the R package ‘junctions’ (Janzen, 2021). By providing the code in an easy to use package, we hope to lower the threshold for other users to apply the theory of junctions to their model system.

## Acknowledgements

We thank Gianni Litti, Molly Schumer, Adriana Suarez-Gonzalez and Quentin Cronk for their help and enthusiasm with respect to our re-analysis of their datasets. We also thank Amaury Lambert who helped building this fruitful collaboration and Arno Siri-Jégousse for a careful reading of a previous version of this manuscript. TJ thanks the Carl von Ossietzky University of Oldenburg for use of their computer cluster CARL and the Center for Information Technology of the University of Groningen for their support and access to the Peregrine high performance computing cluster. VMP was funded by the DGAPA-UNAM postdoctoral program.

## Data Accessibility

We have included the derivations and the numerical solution framework in the R package ‘junctions’, which can be found on CRAN on https://CRAN.R-project.org/package=junctions. All code used in data analysis and visualization for this manuscript can be downloaded from data dryad, from: https://datadryad.org/stash/share/GCJGziFXtI6CV42rl3mcwa92-1nNNMND3PH94QxZL-k (please note that upon publication, this link will be replaced with doi:10.5061/dryad.xwdbrv1c5)

## Author contributions

TJ and VMP jointly designed the research. VMP inferred the ARG based mathematics, TJ verified findings using individual based simulations and analyzed the empirical data. TJ and VMP jointly wrote the paper.

## Supporting information

**S1 Appendix. Sensitivity analysis**. Using individual based simulations, we test how sensitive our new framework is to variation of the different parameters.

**S2 Appendix. Small population size**. We test the validity of our method using a smaller value of the population size (*N* = 1000).

**S3 Appendix. Phasing error**. We extend the analysis done in the main text to the case of a smaller number of markers.

**S4 Appendix. Population size effect on age estimates in empirical data**. We extend the analyses done in the main text for the yeast and swordtail datasets by exploring the impact of population size on the age estimates.

**S5 Appendix. Comparison with ancestry hmm for a larger population size**. We repeat the analysis for Figure 10, but for a population size of *N* = 10, 000.

## Notes

### Competing Interest Statement

The authors have declared no competing interest.

