## Supplementary material for "Estimating the time since admixture from phased and unphased molecular data": S1 Appendix

### S1 Appendix: Sensitivity analysis

Using individual based simulations, we test how sensitive our new framework is to variation in the parameters $N$, $p$ and in the overall rate of recombination $C$ ($C = \sum_{i=1}^{n} d_i$). As in the main text, we simulate with $N = 10,000$, $p = 0.5$ and $C = 1$, we use $n = 10,000$ to mimic coverage found in empirical data (see main text).

#### 1 Population size

Fig 1 shows how sensitive inference is if we infer the time since the onset of admixture using a different value of $N$ as that used to simulate the data ($N = 10,000$). If data from a single chromosome is used and if $N$ is much smaller than the population size used in the simulations (e.g. if $N = 1000$), the tie since admixture tends to be overestimated. In other cases, the inferred time is very similar to the simulated time. In particular, if data from two chromosomes is available (phased or un-phased), population size is of little impact on the inferred time.

#### 2 Recombination rate

Incorrect assessment of recombination rates has drastic effects on the inferred time (2), regardless on the method used to infer the time since admixture. An underestimation of the amount of recombination dramatically inflates the inferred time since admixture, whereas an overestimate of the amount of recombination leads to an underestimation.

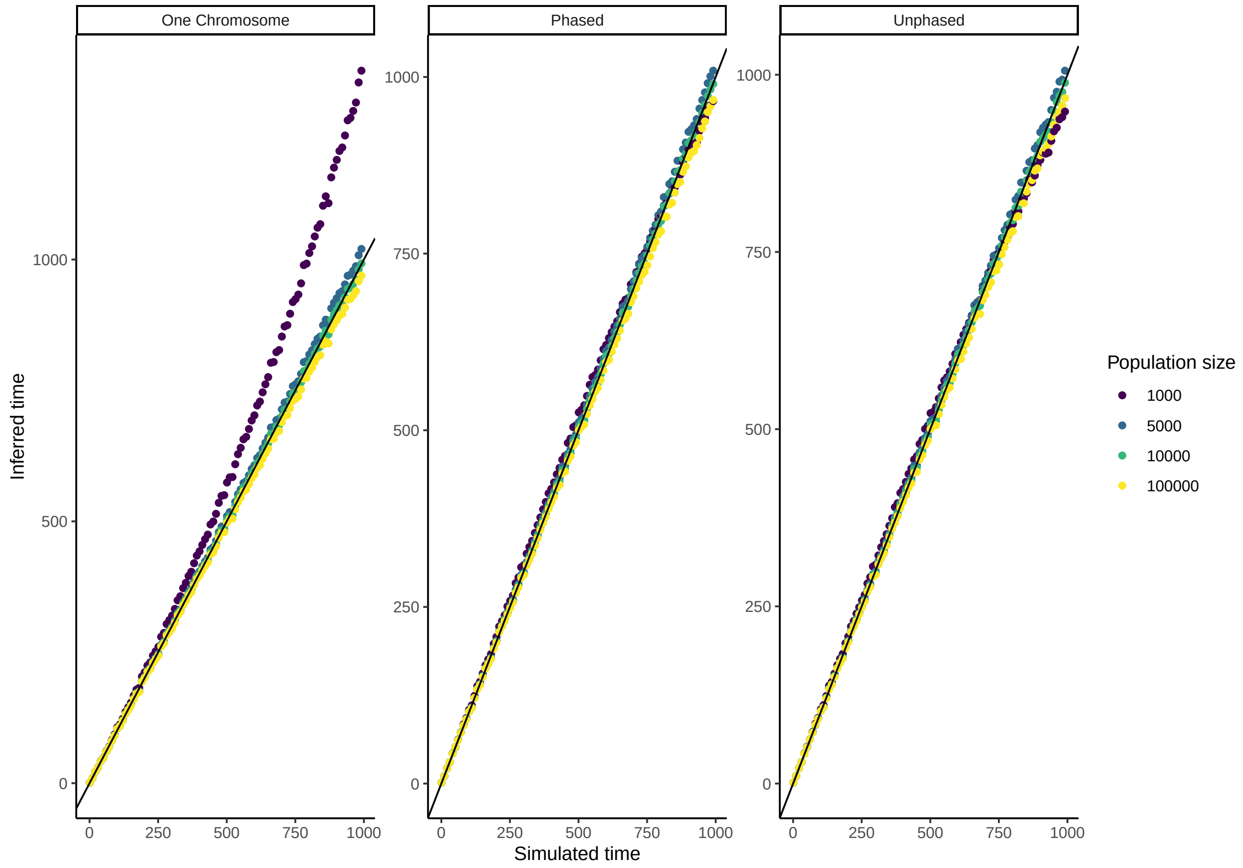


**Fig 1. Sensitivity to $N$.** Estimated time since admixture for data simulated with $N = 10,000$, $p = 0.5$, $C = 1$ and $n = 10,000$. The solid black line indicates the simulated = estimated time, colors indicate the inferred time since admixture assuming a different population size than used to simulate. Shown are the median estimates across 10 replicates, where the time since admixture was inferred for 10 separate individuals per replicate, per time point. The three panels represent the three different methods available: using only information from a single chromosome, using phased information from two chromosomes, or using un-phased information from two chromosomes.

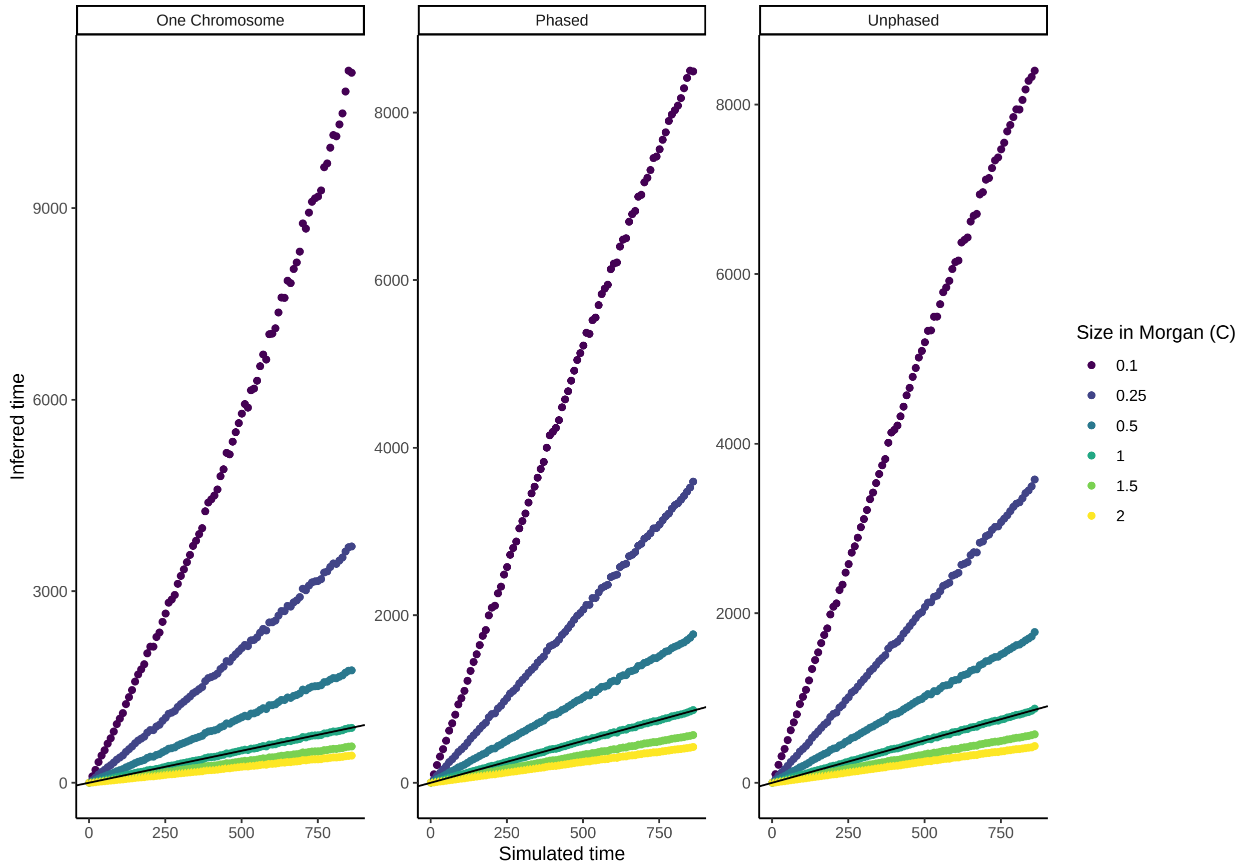


**Fig 2. Sensitivity to $C$.** Estimated time since admixture for data simulated with $N = 10,000$, $p = 0.5$, $C = 1$ and $n = 10,000$. The solid black line indicates the simulated = estimated time, colors indicate the inferred time since admixture assuming different recombination rates than used to simulate. Shown are the median estimates across 10 replicates, where the time since admixture was inferred for 10 separate individuals per replicate, per time point. The three panels represent the three different methods available: using only information from a single chromosome, using phased information from two chromosomes, or using un-phased information from two chromosomes.

### 3 Initial heterozygosity

Incorrect information about the genetic contribution of one of the two source taxa at the onset of hybridization only leads to an overestimate of the time since admixture for extreme deviations from the value used to simulate the data (e.g. only for $p = 0.01$ or 0.99, whilst the data was simulated with $p = 0.5$), if information from both chromosomes is used (phased or un-phased). If only information of a single chromosome is available, incorrect identification of $p$ is more detrimental to the inferred time since admixture, and always leads to an overestimate.

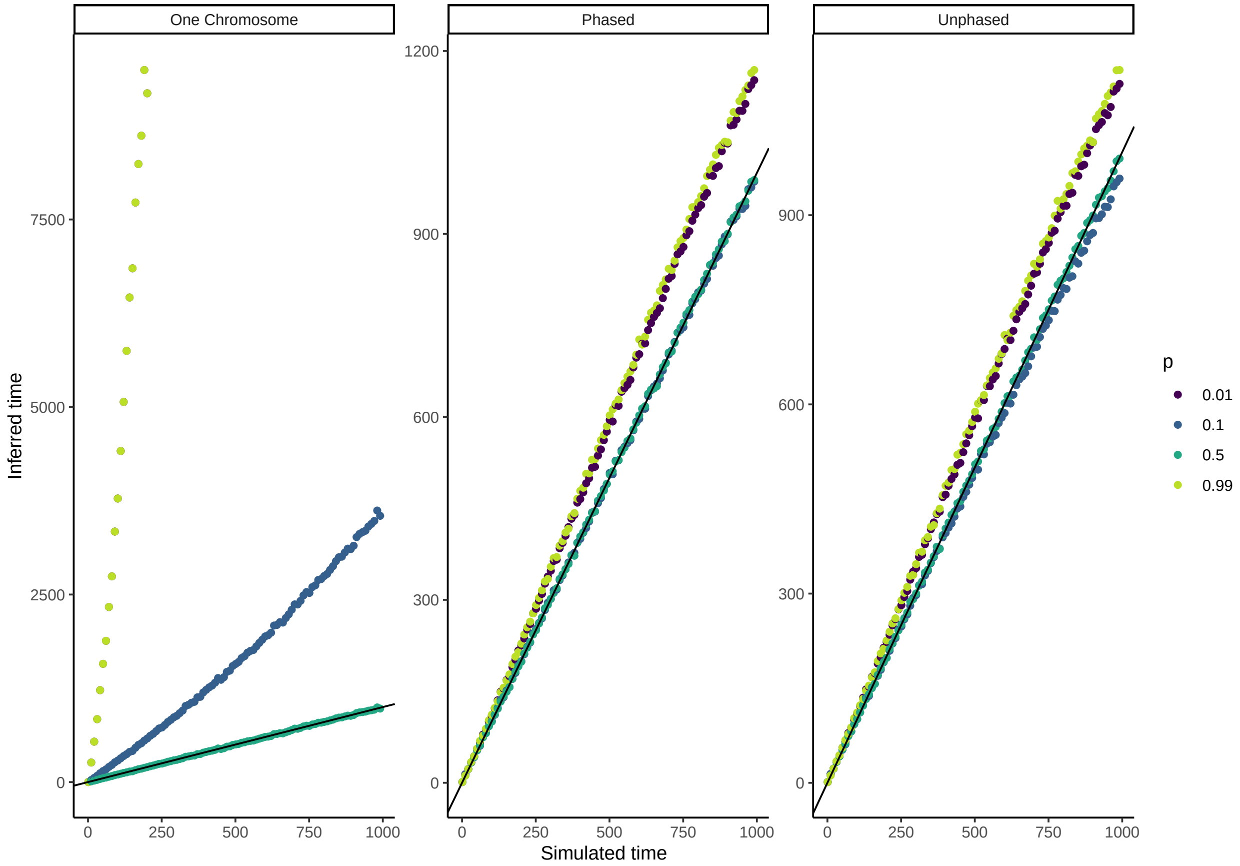


**Fig 3. Sensitivity to $p$.** Estimated time since admixture for data simulated with $N = 10,000$, $p = 0.5$, $C = 1$ and $n = 10,000$. The solid black line indicates the simulated = estimated time, colors indicate the inferred time since admixture assuming different contributions of one of the two source taxa (e.g. $p$) than used to simulate. Shown are the median estimates across 10 replicates, where the time since admixture was inferred for 10 separate individuals per replicate, per time point. The three panels represent the three different methods available: using only information from a single chromosome, using phased information from two chromosomes, or using un-phased information from two chromosomes.

### 4 Number of markers

Having a low number of markers drastically increases the uncertainty of the estimated time since admixture, but only for very low numbers (e.g. <1000 markers per chromosome) (Fig 4). Although the median estimates remain identical to the expected time, the variance becomes very high when the number of markers is very low.

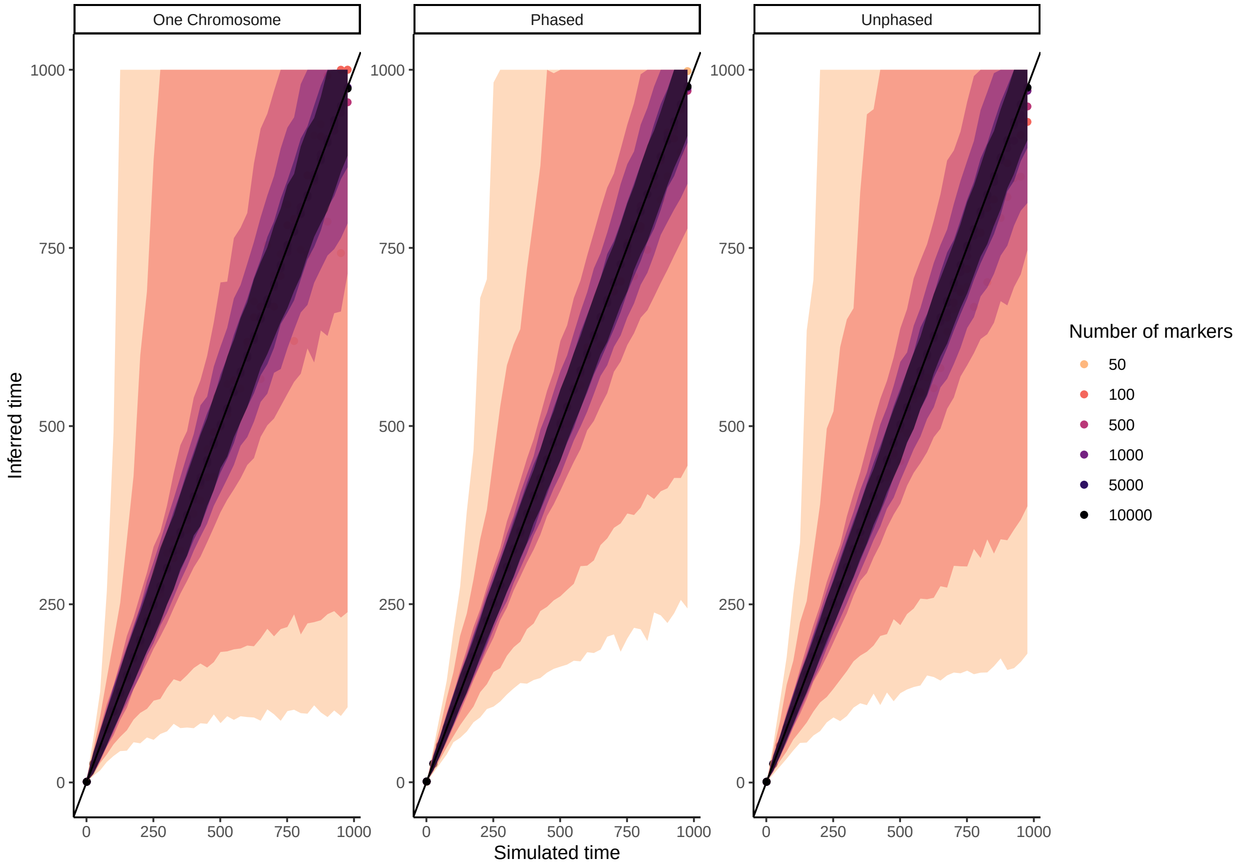


**Fig 4. Sensitivity to $n$.** Estimated time since admixture for data simulated with $N = 10,000$, $p = 0.5$ and $C = 1$. The solid black line indicates the simulated = estimated time, colors indicate the inferred time since admixture assuming different numbers of markers. Shown are the median estimates across 100 replicates, and the 95% envelope. The time since admixture was inferred for 10 separate individuals per replicate, per time point. The three panels represent the three different methods available: using only information from a single chromosome, using phased information from two chromosomes, or using un-phased information from two chromosomes.
