## Supplementary material for "Estimating the time since admixture from phased and unphased molecular data": S2 Appendix

### S2 Appendix: Small population size

In this appendix, we test the validity of our maximum likelihood approach and its sensitivity to the different parameters, but using a smaller value of the population size in the simulations ($N = 1000$ instead of 10000).

#### 1 Validation of the method

In this section, we use the same parameters as in the individual based simulations in the main text but with a smaller population size ($N = 1000$).

In Fig 1 we compare the methods we have developed here to previous methods based on the theory of junctions. Again, we observe that, when the number of markers is low, previous methods, that do not take into account marker spacing, tend to underestimate the time since admixture, which is not the case for our methods.

Fig 2 shows that using the one chromosome method only small hybridization times can accurately be inferred. When the hybridization times are longer than a threshold value (around 5000 generations for $H_0 = 0.5$), the method tends to overestimate the time since admixture. This is due to the fact that there is a plateau in the function giving the number of junctions per unit of time (equation (2) in the main text). This phenomenon is called the maximum packing density of junctions.

However, with the method using information from two homologous chromosome, this problem did not appear and we could always provide an accurate estimation of the time since admixture. The same problem would have appeared if we had performed longer simulations. However, hybridization occurs in short timescales and longer timescales are incompatible with the hypothesis that we can ignore mutation events.

Fig 3 shows a comparison between the two methods that use information from two homologous chromosomes, whether the data is phased or unphased. For short times (below 5000 generations), the phased method provides slightly better results than the unphased method (the relative error is smaller). However, for long times, both methods perform equally well.

### 2 Sensitivity analysis

We also tested if changing the population size to $N = 1000$ changes the sensitivity of our method to the different parameters.

#### 2.1 Population size

Varying the population size has some interesting effects (see Fig 1 in S1 Appendix). When using information from a single chromosome we observe that if $N$ is smaller than the population size in the simulations, the time since admixture is over estimated while, if $N$ is larger it is underestimated. When using information from two chromosomes, whether it is phased or unphased, overestimating the population size initially increases the age estimate, although extreme overestimates contrastingly lead to an underestimate.

#### 2.2 Recombination rate

If we vary the amount of recombination (Fig 5), we observe that higher levels of recombination (e.g. overestimates of recombination) lead to a younger age. When using information from two chromosomes, this changes the age estimate with a constant amount. But when restricted to a single chromosome, an increased recombination leads to levelling off of the age estimate due to reaching age horizon resulting from the maximum packing density of junctions sooner.

#### 2.3 Initial heterozygosity

We find very similar results S1 Appendix. Fig 6 shows that varying the initial heterozygosity has a strong impact in the method that uses one chromosome. However, when using information from the two homologous chromosomes, errors in the estimation of $p$ don't yield such important errors in the estimation of the time since admixture.

#### 2.4 Number of markers

With a smaller population size, we are able to simulate to a much larger number of generations and show that the accuracy of inference tends to be retained, provided that the number of markers is large (5,000 - 10,000 markers per chromosome), and that information on both chromosomes is used (Fig 7). When information of only a single chromosome is used, accuracy in admixture time inference is low. When information from both chromosomes is used, we observe that when the admixture time is above a certain threshold, that depends on the number of markers, inference becomes impossible (the 95% enveloppe becomes extramely large and biased towards high values).

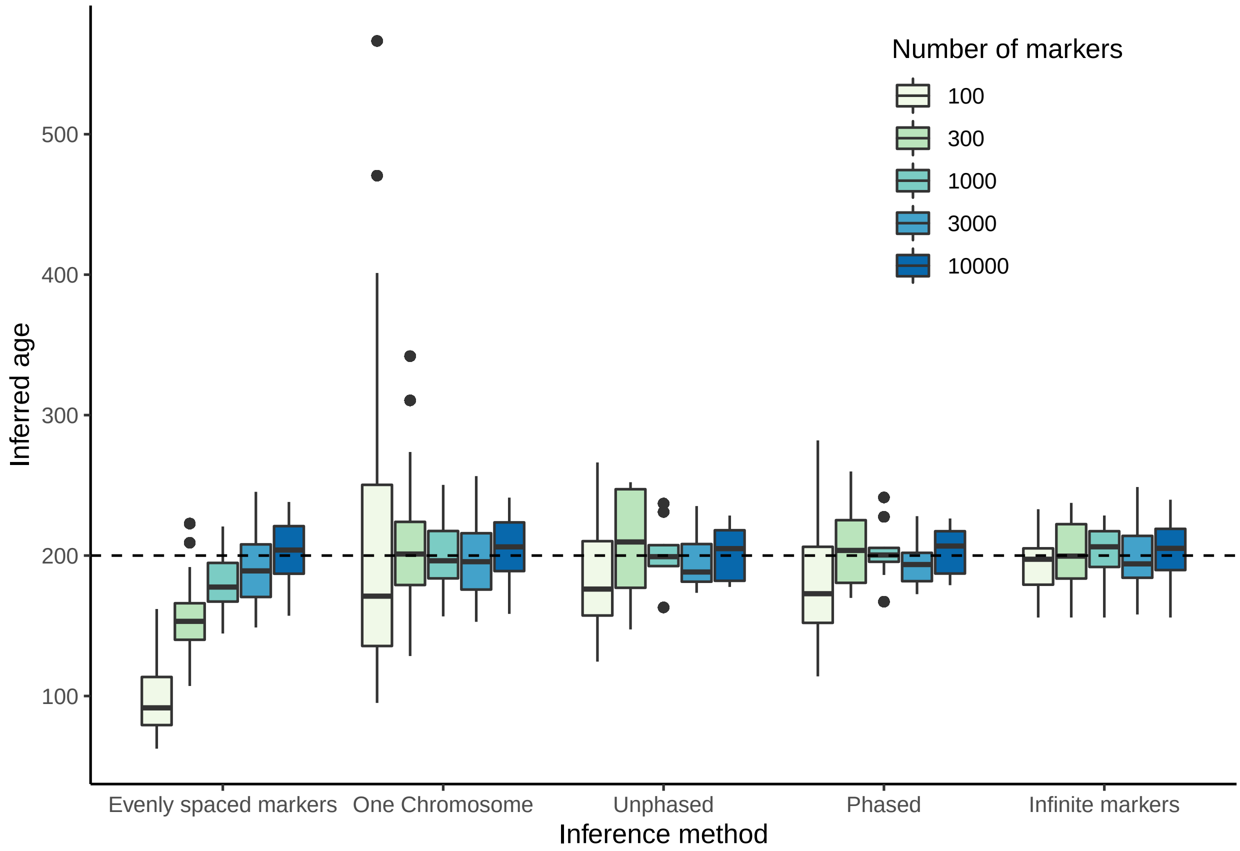


**Fig 1. Inferred time using different methods.** Shown are the median estimates (dots) for 100 replicates, where in each replicate 10 individuals were analyzed. The dashed line indicates the simulated time. 'Evenly spaced markers' corresponds to the method in Janzen et al. (2018). 'Infinite markers' corresponds to an idealized scenario where ancestry is known for every locus in the chromosome and is there to quantify the amount of randomness in the process. The population size was 1,000 individuals, and 10,000 randomly spaced markers were used.

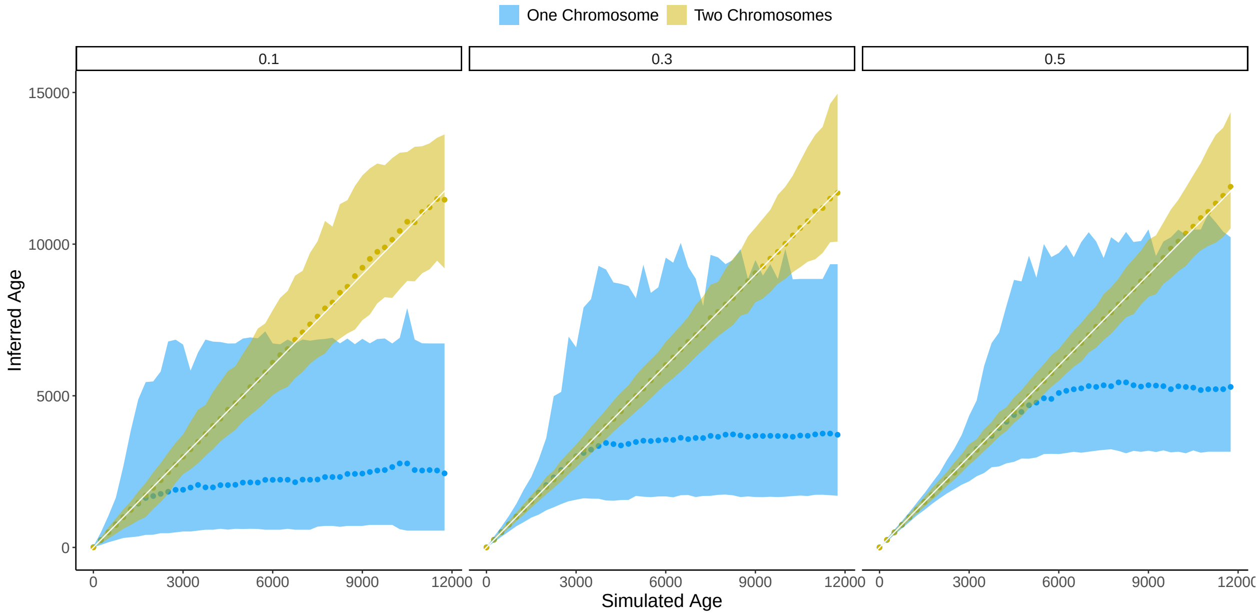


**Fig 2. Inferred time versus simulated time using one or two chromosomes.** Shown are the median estimates (dots) for 100 replicates, where in each replicate 10 individuals were analyzed. The solid white line indicates the observed is equal to expected line and the shaded area indicates the 95% percentile. Shown are results using junction information from one chromosome (blue) and results using information from two chromosomes (gold). Numbers above the plots indicate the initial heterozygosity. The population size was 1,000 individuals, and 10,000 randomly spaced markers were used.

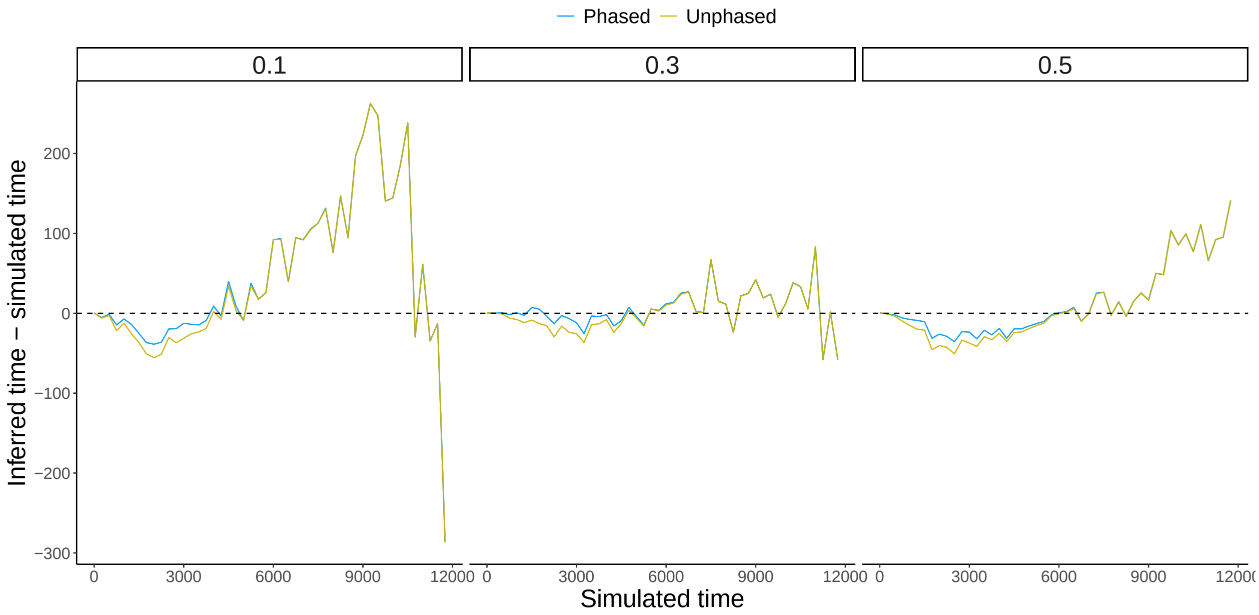


**Fig 3. Accuracy in age estimate using the unphased framework versus the phased framework.** Shown is the median difference across 100 replicates. Shown are results for three different initial heterozygozities, indicated at the top of each plot. The population size was 1,000 individuals, and 10,000 randomly spaced markers were used.

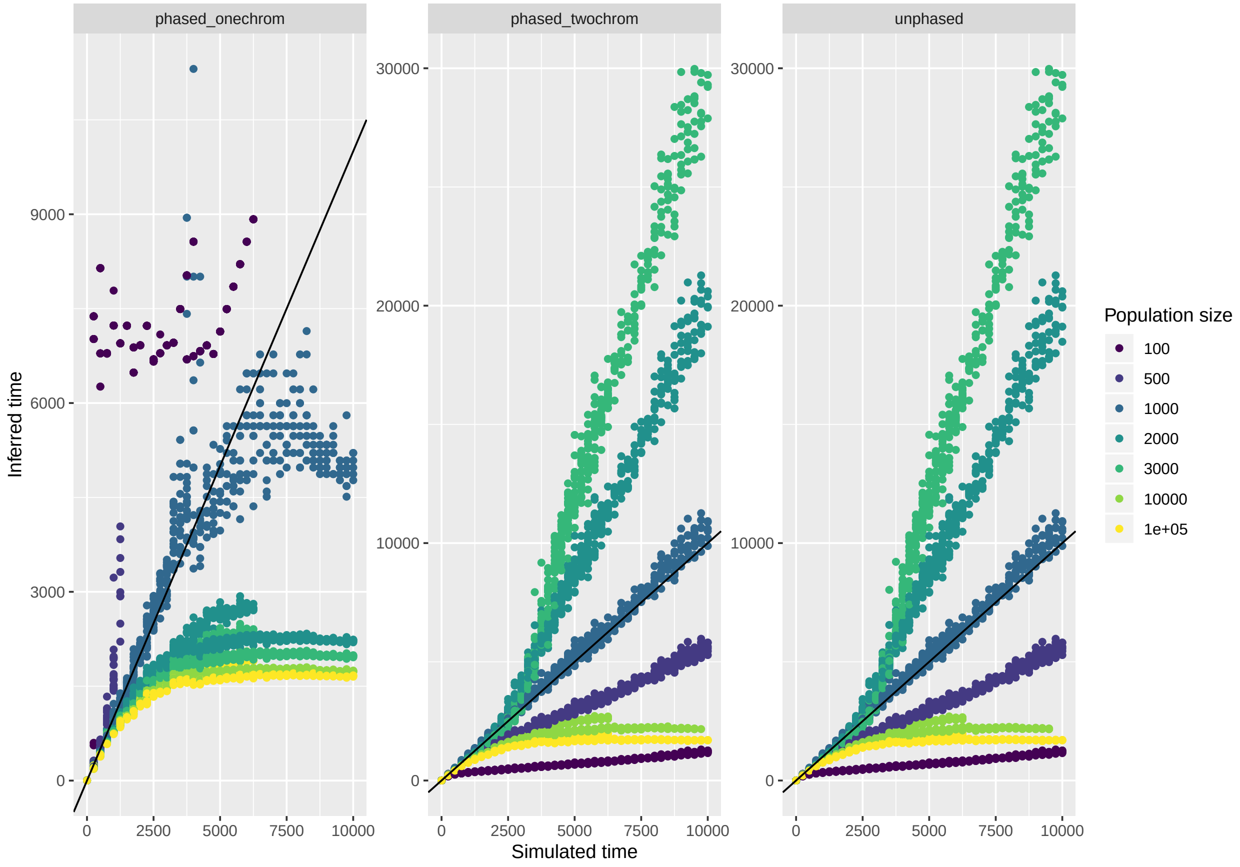


**Fig 4. Sensitivity to $N$** Estimated time since admixture for data simulated with $N = 1000$, $p = 0.5$ and $C = 1$. The solid black line indicates the simulated = estimated time, colors indicate the inferred time since admixture assuming a different value of $N$ than used to simulate..

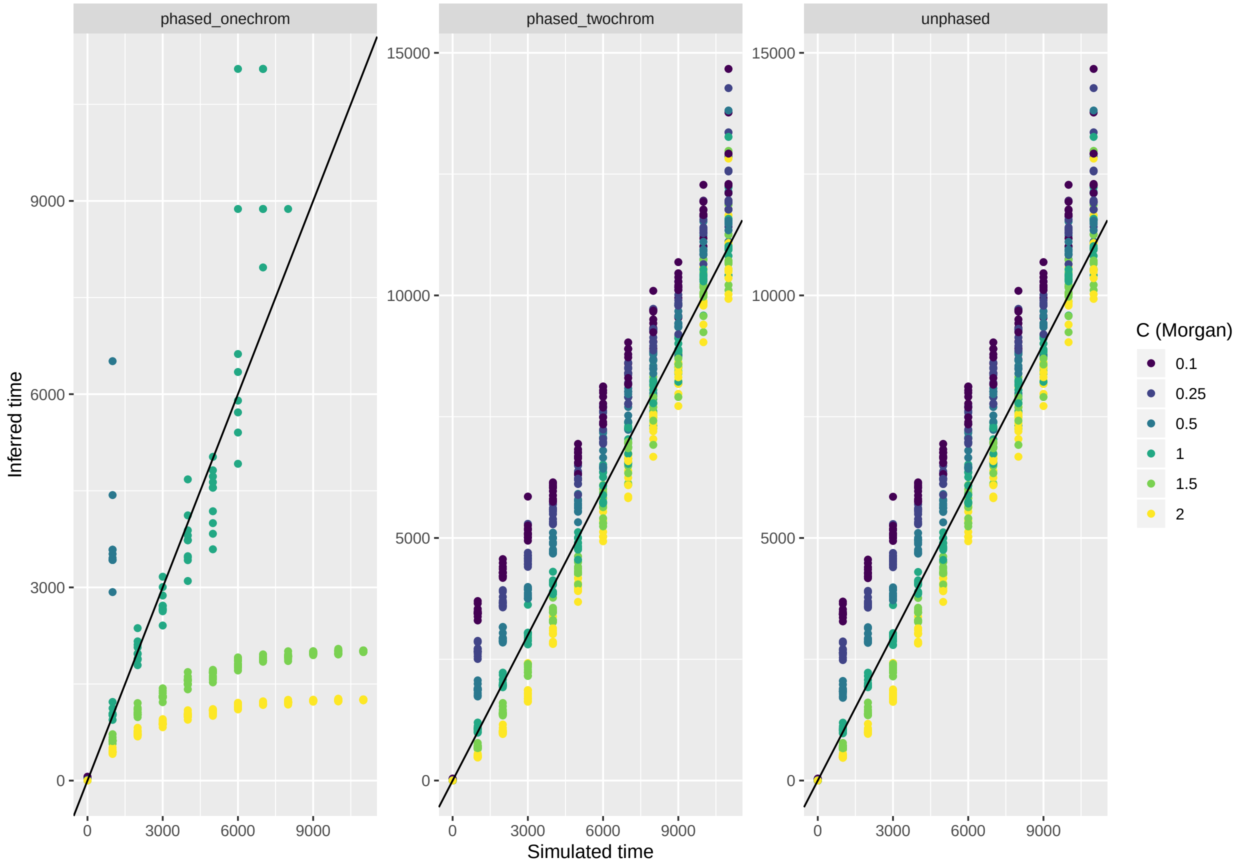


**Fig 5. Sensitivity to $C$** Estimated time since admixture for data simulated with $N = 1000$, $p = 0.5$ and $C = 1$. The solid black line indicates the simulated = estimated time, colors indicate the inferred time since admixture assuming a different rate of recombination than used to simulate.

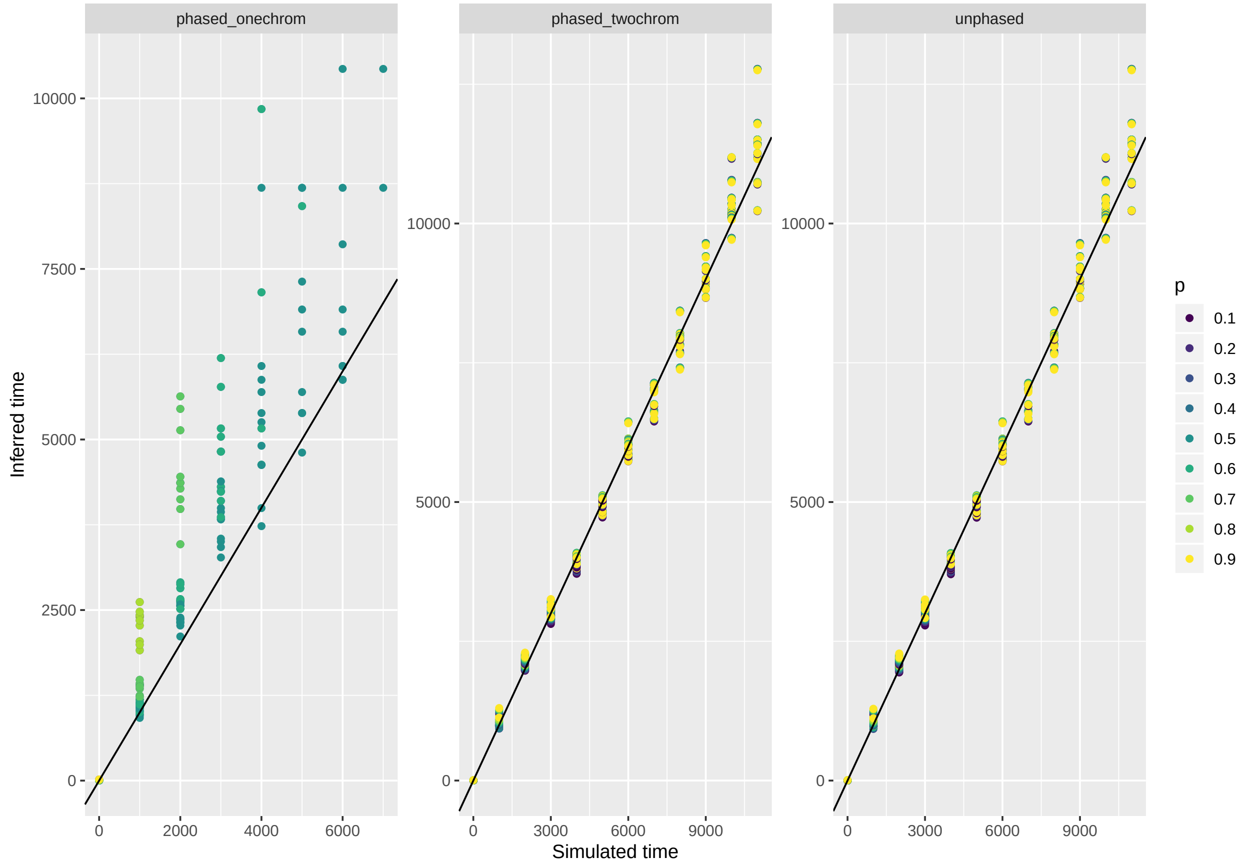


**Fig 6. Sensitivity to $p$.** Estimated time since admixture for data simulated with $N = 1000$, $p = 0.5$ and $C = 1$. The solid black line indicates the simulated = estimated time, colors indicate the inferred time since admixture assuming a different value of $p$ than used to simulate.

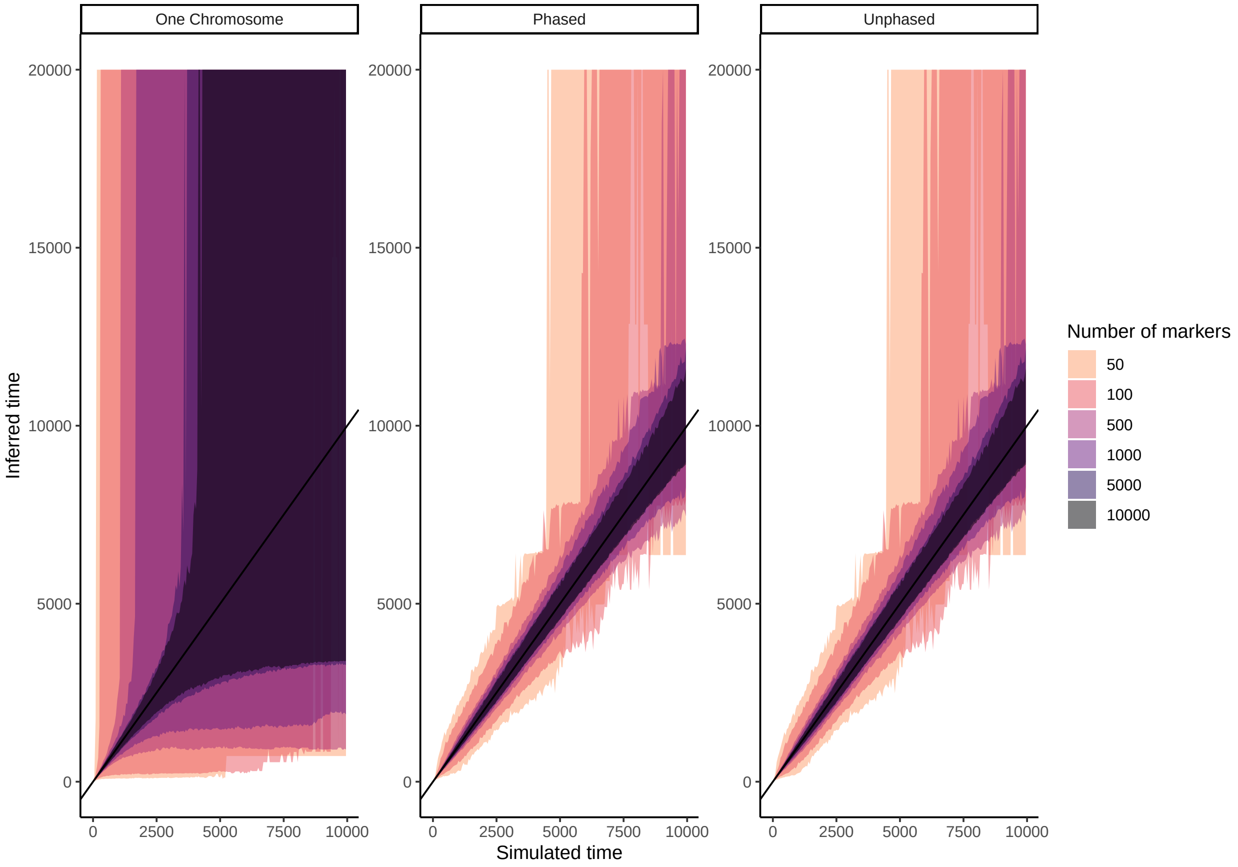


**Fig 7. Sensitivity to $n$.** Estimated time since admixture for data simulated with $N = 1,000$, $p = 0.5$ and $C = 1$. The solid black line indicates the simulated = estimated time, colors indicate the inferred time since admixture assuming different numbers of markers. Shown are the median estimates across 100 replicates, and the 95% envelope. The time since admixture was inferred for 10 separate individuals per replicate, per time point. The three panels represent the three different methods available: using only information from a singleinformation from two chromosomes, or using unphased information from two chromosomes.
