## Supplementary material for "Estimating the time since admixture from phased and unphased molecular data": S3 Appendix

### S3 Appendix: Phasing error

We extend the analysis done in the main text by an analysis where we explore the error in inference of admixture time using only 500 or 1,000 markers. In the main text we only focused on 10,000 markers, which is much larger than the simulated time. In this scenario, incorrectly phased markers *always* introduce a fake junction, but never remove a junction. In contrast, if the number of junctions is lower or equal to the simulated time, an incorrectly phased marker might also remove an observed junction. Instead of focusing on the same percentage of phasing error, we have chosen to use the same number of incorrectly phased markers - in order to avoid having no incorrectly phased markers at all (0.25% of 500 markers has an expected value lower than 1). We simulate again with a population of 10,000 individuals, for $p = 0.5$ and $C = 1$. We find that including of incorrectly phased markers tends to increase the age estimate, even when the number of markers is relatively low. Please note that the error might seem incredibly large, but this might be due to the fact that the absolute number of incorrect markers is high compared to the total number of markers.

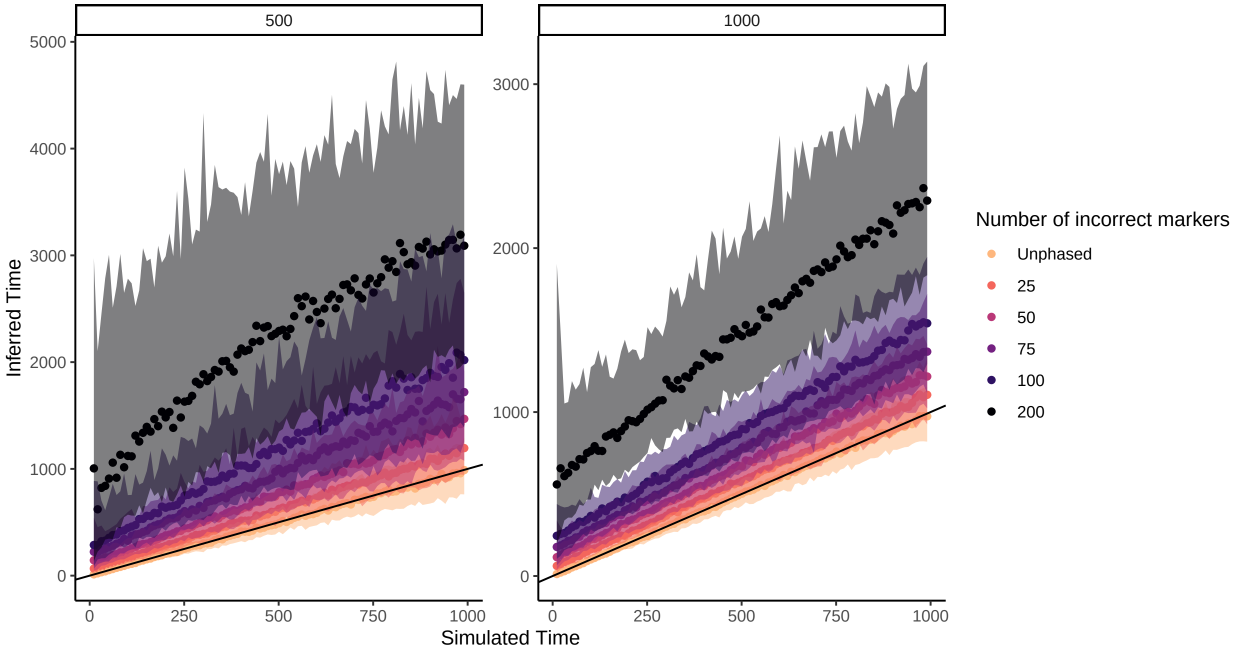


**Fig 1. Effect of phasing error on the estimated time since admixture.** Estimated time since admixture for data simulated with $N = 10,000$, $p = 0.5$ and $C = 1$. The solid black line indicates the simulated = estimated time, colors indicate the inferred time since admixture including a varying number of incorrectly phased markers. The colors indicate the 95% envelope. The left panel shows results for $n = 500$, the right panel shows results for $n = 1000$.
