## Supplementary material for "Estimating the time since admixture from phased and unphased molecular data": S4 Appendix

### S4 Appendix: Population Size in empirical datasets

#### 1 Yeast

We have repeated the analysis in the main text, but varied the population size in [10, 100, 1000, 10000, 10000]. The results show (Fig 1) that with the age estimate is irrespective of population size, as long as the population size is above 100 individuals. Only for very small population sizes (10 individuals), does the age estimate increase, which brings the age estimates obtained with the two older recombination rate estimates (Cherry et al., 1997; Mancera et al., 2008) closer to the number of generations used in the experiment.

#### 2 Swordtail fish

We repeat the analysis shown in the main text, but with varying population size. Whereas the main text used the population size estimate of 1830 individuals as obtained from Schumer et al. (2014), here we explore the sensitivity of the age estimate to varying population size, by varying the value used for $N$ in [5000, 10000, 100000, 1000000]. We find that using a (much) larger population size only reduces the age estimate to a small extent.

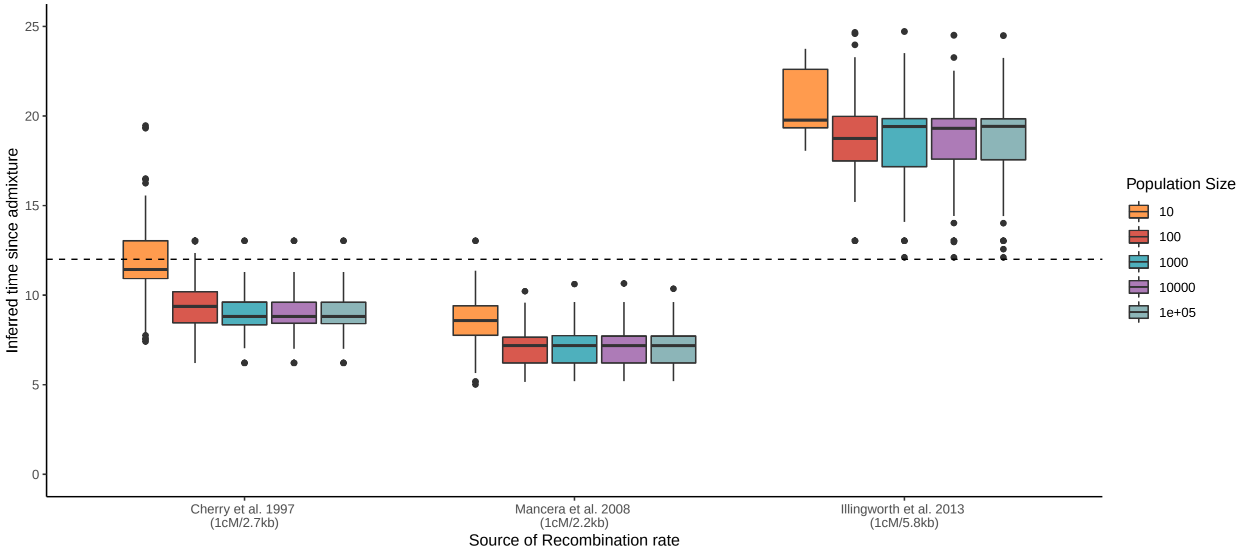


**Fig 1. Inferred age for F12 Hybrid Yeast (*Saccharomyces cerevisiae*) individuals.** (A) Inferred age for three different recombination rates: $1cM/2.7kb$ (Cherry et al., 1997), $1cM/2.2lb$ (Mancera et al., 2008) and $1cM/5.8kb$ (Illingworth et al., 2013). Shown is the distribution of inferred ages across 171 individuals, inferred using different population sizes.

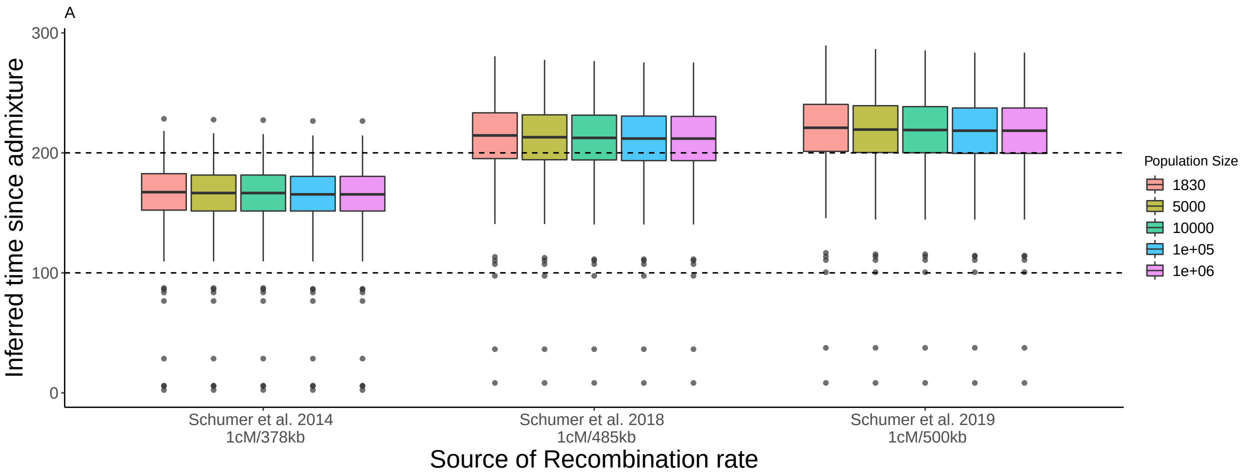


**Fig 2. Inferred age for hybrid *Xiphophorus* fish from Tlatemaco (Mexico)**. Shown are age inferences based on three different recombination maps: $1cM/370kb$ (Schumer et al., 2014), $1cM/485kb$ (Schumer et al., 2018) and $1cM/500kb$ (Powell et al., 2020). Dotted lines indicate the hypothesized age limits of the population. Colors indicate different population sizes used during the age inference.

E. Mancera, R. Bourgon, A. Brozzi, W. Huber, and L.M. Steinmetz. High-resolution mapping of meiotic crossovers and non-crossovers in yeast. *Nature*, 454(7203):479–485, 2008.

C.J.R. Illingworth, L. Parts, A. Bergström, G. Liti, and V. Mustonen. Inferring genome-wide recombination landscapes from advanced intercross lines: application to yeast crosses. *PLoS One*, 8(5):e62266, 2013.

M. Schumer, R. Cui, D. L. Powell, R. Dresner, et al. High-resolution mapping reveals hundreds of genetic incompatibilities in hybridizing fish species. *Elife*, 3:e02535, 2014.

M. Schumer, C. Xu, D. L. Powell, A. Durvasula, et al. Natural selection interacts with recombination to shape the evolution of hybrid genomes. *Science*, 360(6389):656–660, 2018.

D. L. Powell, M. García-Olazábal, M. Keegan, P. Reilly, et al. Natural hybridization reveals incompatible alleles that cause melanoma in swordtail fish. *Science*, 368(6492):731–736, 2020.
