## Supplementary material for "Estimating the time since admixture from phased and unphased molecular data": S5 Appendix

### S5 Appendix: Comparison with ancestry hmm for larger population sizes

In the main text, figure 10 shows inference of the time since admixture for markers of varying quality, using $N = 1000$. Here, we repeat the analysis, but for $N = 10000$. Figure 1 shows that ancestry hmm is more accurate in inferring the time since admixture when the population size is large, and that the differences between the inferred age by ANCESTRY HMM and our method converge towards the same estimate as the number of markers increases.

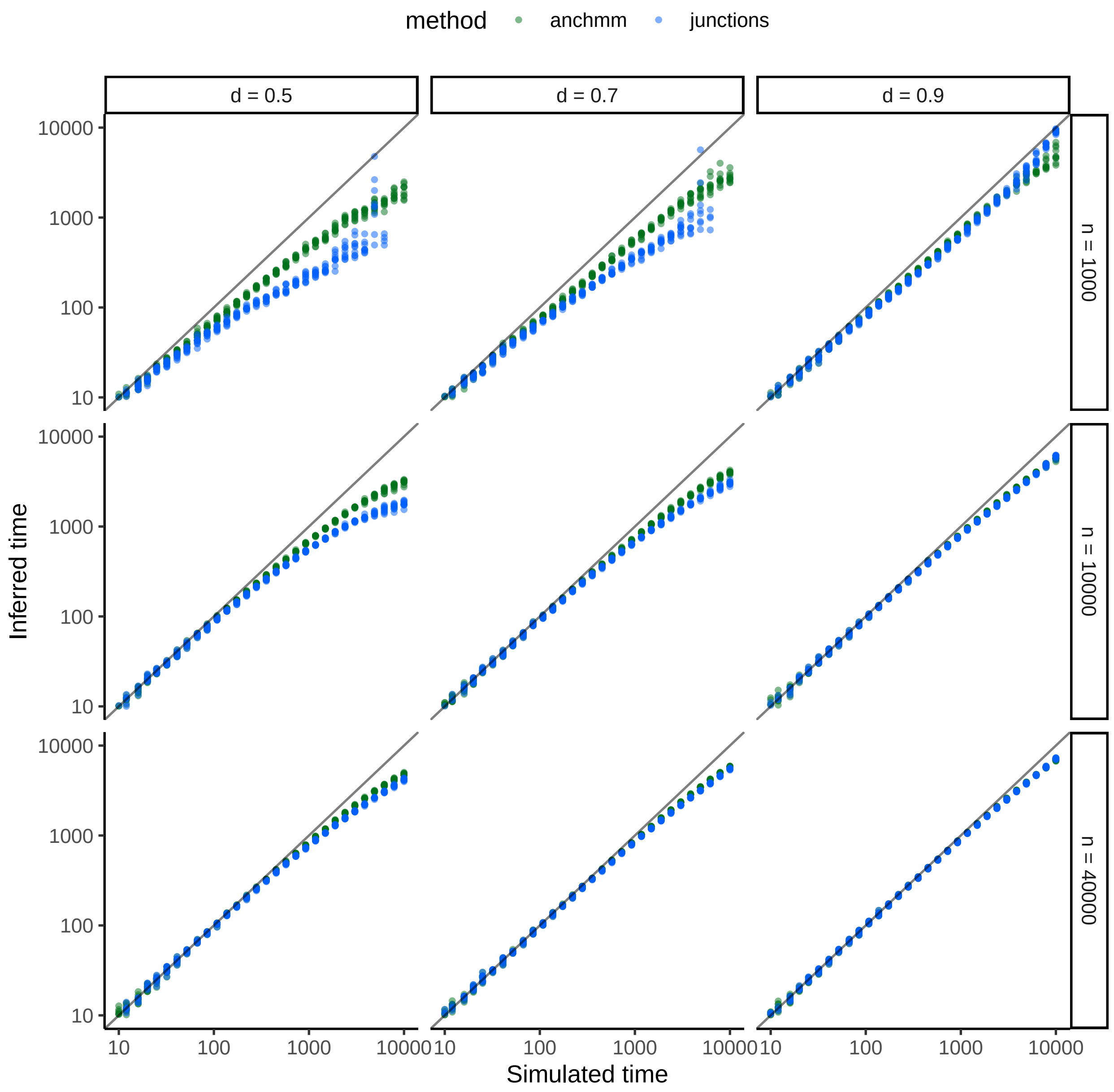


**Fig 1. Comparison in estimating the time since admixture between ancestry hmm and the method proposed here, for a large population of $N = 10000$.**. The solid black line indicates the simulated = estimated time. Dots indicate the inferred ages, with the green dots representing ages inferred by ANCESTRY HMM and the blue dots indicate ages inferred by the junctions framework. Age estimates are based on simulated data with uncertain ancestry, where uncertainty in ancestry is reflected by the allele frequency differential (Shriver et al., 1997). Because the method proposed here does not include ancestry uncertainy, local ancestry as inferred by ANCESTRY HMM was used.
